# Comparative single-cell transcriptomic atlases reveal conserved and divergent features of drosophilid central brains

**DOI:** 10.1101/2023.11.22.568274

**Authors:** Daehan Lee, Richard Benton

**Affiliations:** Center for Integrative Genomics Faculty of Biology and Medicine University of Lausanne CH-1015, Lausanne Switzerland; Department of Biological Sciences College of Natural Sciences Sungkyunkwan University, Suwon 16419 Republic of Korea

## Abstract

To explore how brains change upon species evolution, we generated single-cell transcriptomic atlases of the central brains of three closely-related but ecologically-distinct drosophilids: the generalists *Drosophila melanogaster* and *Drosophila simulans*, and the noni fruit specialist *Drosophila sechellia*. The global cellular composition of these species’ central brains is well-conserved, but we predicted a few cell types (perineurial glia, sNPF and Dh44 peptidergic neurons) with divergent frequencies. Gene expression analysis revealed that distinct cell types within the central brain evolve at different rates and patterns; notably, glial cell types exhibit the greatest divergence between species. Compared to *D. melanogaster*, the cellular composition and gene expression patterns of the central brain in *D. sechellia* display greater deviation than those of *D. simulans* - despite their similar phylogenetic distance from *D. melanogaster* - that the distinctive ecological specialization of *D. sechellia* is reflected in the structure and function of its brain. Expression changes in *D. sechellia* encompass metabolic and ecdysone signaling genes, suggestive of adaptations to its novel ecological demands. Additional single-cell transcriptomic analysis on *D. sechellia* revealed genes and cell types responsive to dietary supplement with noni, pointing to glia as sites for both physiological and genetic adaptation to novel conditions. Our atlases represent the first comparative analyses of “whole” central brains, and provide a comprehensive foundation for studying the evolvability of nervous systems in a well-defined phylogenetic and ecological framework.

## Introduction

Animal nervous systems contain hundreds to billions of cells. These cells, encompassing neurons and glia, can be categorized into a large number of different types – based upon developmental, anatomical, molecular and functional properties^1,2^ – with diverse roles. For example, neurons act in sensory detection, information processing and locomotor control, while glia generally have support functions, including as structural/insulating scaffolds, in nutrient supply and through removal of cellular debris and toxins^3,4^. The complement of cells in the nervous system of an extant species arises from ongoing evolutionary processes, where external selection pressures can lead to emergence of new (or modified) cell types that fulfill functions conferring a fitness advantage (e.g., detecting a novel pertinent sensory stimulus, fine-tuning a motor action, or for modulating energy homeostasis). Conversely, cells whose function no longer contributes to organismal fitness might be lost (or repurposed). Understanding the genetic mechanism and selective pressures underlying the gain, loss and modification of cell types can reveal how and why nervous systems change over evolutionary timescales, as well as basic insights into the development and function of neural circuits^5,6^. Until recently, our understanding of nervous system evolution relied heavily on correlations of differences in the macroscopic (neuro)anatomy and behavior of different species^5–8^, limiting our mechanistic (i.e., genetic) understanding of how evolutionary changes occur. Single-cell transcriptomic approaches have enormous potential to advance this knowledge, by enabling cataloging of neurons and glia and their molecular relationships in various species to suggest hypotheses for how – and why – divergence in cellular composition has occurred. For example, profiles of cerebellar output neurons in mice, chickens and humans suggested that developmental duplications of subsets of these cells underlie the large expansion of the human cerebellum^9^, while comparisons of cell type-specific transcriptomes in reptiles, amphibians and vertebrates have provided insights into both ancient cell types and mammalian-specific innovations of the cortex and other brain regions^10–12^. Such surveys can also relate gene and cell type evolution, such as a comparison of hypothalamic neuronal populations in fish, which revealed that species-specific cell types were often characterized by expression of species-specific gene paralogs^13^.

A limitation of studying nervous system evolution in vertebrates is the very large number of cells in their brains: a sampling of >3 million cells from diverse sites in the human brain is a minuscule fraction of the estimated 100-200 billion cells of this organ^14^, while a “whole”-brain scRNA-seq atlas of the mouse *Mus musculus* profiled ∼7 million neurons^15^, but this still represents only about 10% of the total^16^. Moreover, relating structural differences to ecologically-relevant functional differences is often challenging. In this context, flies of the *Drosophila* genus define an excellent model clade for investigating nervous system evolution for several reasons. First, these species have relatively compact central brains, comprising ∼43,000 cells (of which ∼90% are neurons) in *Drosophila melanogaster*^17^. Second, *D. melanogaster* has been the focus of a wealth of molecular, anatomical and functional studies relating brain structure to function over several decades^18–20^, including the generation of single-cell transcriptomic atlases^21–23^. Third, different drosophilid species have adapted to diverse ecological niches – with different food sources, climate conditions, competitors, pathogens and predators – and display numerous species-specific behaviors, including sensory responses to environmental stimuli and motor actions, such as courtship song production^24,25^.

Amongst drosophilids, the trio of *D. melanogaster*, *Drosophila simulans* and *Drosophila sechellia* present a particularly interesting set of species for comparative neurobiology. These species diverged from a common ancestor 3-5 million years ago, with *D. simulans* and *D. sechellia* diverging much more recently (100-250,000 years ago)^26,27^ (Fig. 1a). While *D. melanogaster* and *D. simulans* are cosmopolitan generalists, with overlapping geographic ranges and similar broad use of fermenting vegetal substrates for feeding and breeding, *D. sechellia* is endemic to the Seychelles archipelago and has evolved extreme ecological specialization, spending most or all of its life cycle exclusively on the “noni” fruit of the shrub *Morinda citrifolia*^28^. As noni fruit is toxic for other drosophilids, including *D. simulans* inhabiting the Seychelles^29,30^, this niche specialization might alleviate interspecific competition, and possibly essential nutritional benefits^31^. Commensurate with its unique ecology, *D. sechellia* displays many behaviors that are distinct from its generalist relatives, including olfactory and gustatory preferences, circadian plasticity and certain reproductive behaviors^32–37^. Some of these behaviors have been linked to structural and/or functional changes in the peripheral nervous system. For example, several populations of olfactory sensory neurons display increased sensitivity to noni-derived odors in *D. sechellia* compared to *D. melanogaster* and *D. simulans*, due to coding mutations in specific olfactory receptors. Furthermore, a number of olfactory sensory neuron populations are several-fold larger (or smaller) in *D. sechellia*, presumably due to changes in the developmental patterning of the olfactory organs^32,33,38–40^.

**Figure 1.**
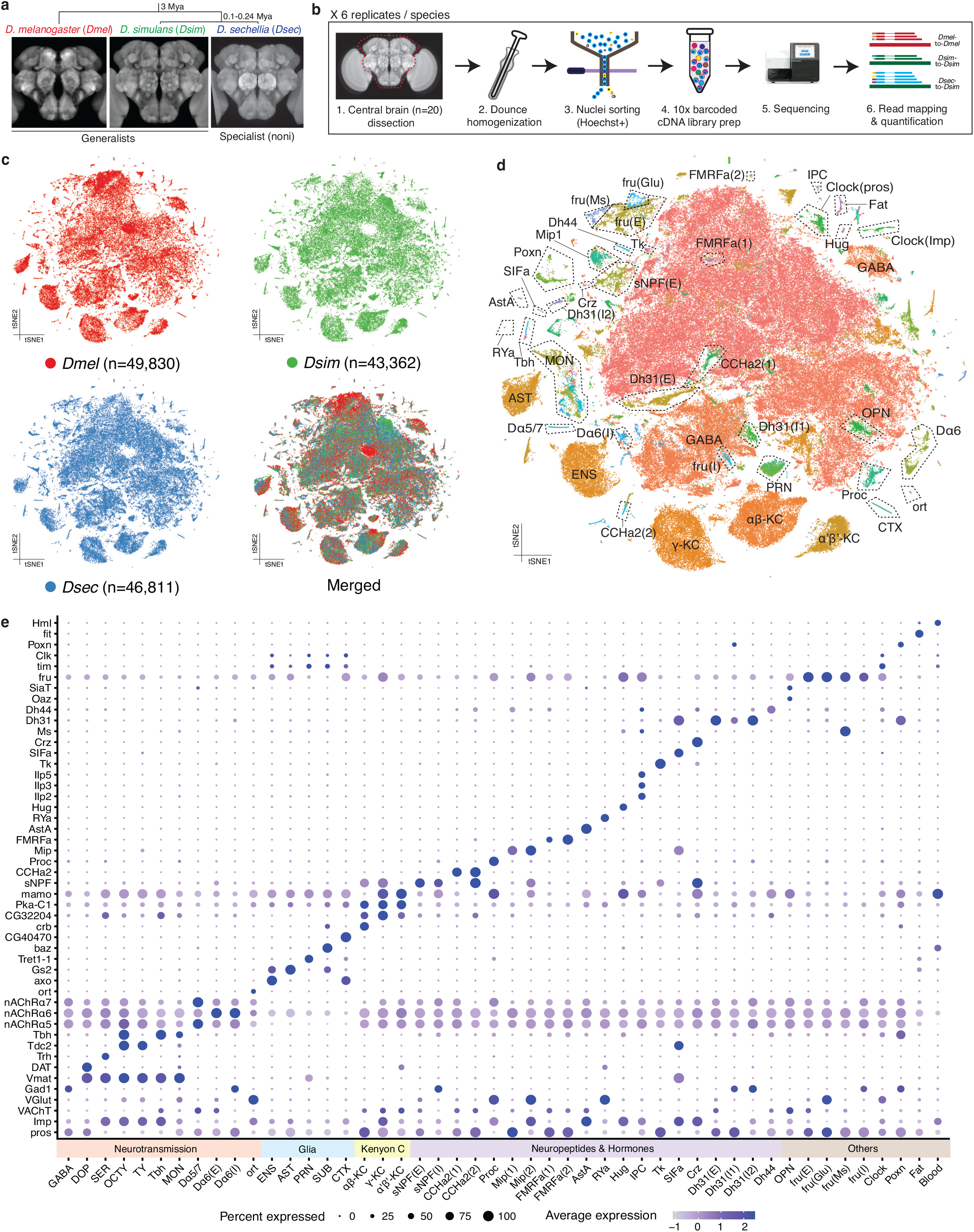
Integrated single-cell transcriptomic atlas of *D. melanogaster*, *D. simulans* and *D. sechellia*. (a) Top: phylogeny of the drosophilid species studied in this work. Bottom: images of reference central brains for these species (all female; source: *D. melanogaster*^71^; *D. simulans*^72^; *D. sechellia*^73^. (b) Workflow of the single-nucleus RNA-sequencing of drosophilid central brains. (c) tSNE plots of *D. melanogaster* (red), *D. simulans* (green) and *D. sechellia* (blue) central brain cells from an integrated dataset after RPCA integration. In the bottom right plot, all cells from the three species are merged. (d) tSNE plot of the integrated and annotated datasets. Cells are colored by the SNN-based clustering and subclustering analyses. (e) Dot plot summarizing the expression of genes used for the annotation of 48 cell types. The “Blood” cluster might correspond to a small number of hemocytes that cross the blood-brain barrier from the hemolymph^74^.

Despite these advances in defining causal, or at least correlative, relationships between peripheral sensory neuron changes and species-specific behaviors and ecologies of these drosophilids, we know essentially nothing about if and how their central brains have evolved. Here, we performed comparative single-cell transcriptomic analyses of the central brains of this drosophilid trio to produce, to our knowledge, the first comparative whole central brain atlases in any species. Our results reveal conserved and divergent features of the cellular composition and gene expression patterns in these drosophilid’ central brains as well as signals and patterns of brain evolution upon niche specialization. This study provides a valuable dataset for future studies to investigate the genetic and cellular basis of brain evolution.

## Results

### Identification of diverse conserved cell types in drosophilid central brains

We generated central brain comparative single-cell transcriptomic atlases of *D. melanogaster*, *D. simulans* and *D. sechellia* through single-nucleus RNA-sequencing (snRNA-seq) (Fig. 1a,b, Methods). In brief, we dissected the brains of 5-day-old, mated female adults cultured on standard food medium, and removed the optic lobes. All steps – tissue dissection, nuclear isolation, RNA extraction, library preparation and sequencing – were performed in parallel for the three species for six biological replicates, each consisting of 20 brains per species. During the processing of snRNA-seq data with the Cell Ranger software (ver. 6.0.1)^41^, sequence reads from *D. melanogaster* and *D. simulans* were mapped to their respective genomes. However, due to the suboptimal assembly and annotation quality of the *D. sechellia* reference genome, we aligned *D. sechellia* sequence reads to the *D. simulans* genome (Methods). In total, the number of nuclei sequenced and analyzed per species (*D. melanogaster* = 49,830; *D. simulans* = 43,362; *D. sechellia* = 46,811) all exceed the estimated cell number in the central brain of *D. melanogaster* (∼43,000^17^). Our transcriptomic atlases therefore comprise ∼1X cell coverage of their central brains. On average, we detected the expression of 888, 816 and 778 genes per cell for *D. melanogaster*, *D. simulans* and *D. sechellia*, respectively, corresponding to 1,766, 1,487 and 1,377 Unique Molecular Identifiers. Detailed metrics for the snRNA-seq results can be found in Supplementary Table 1.

To integrate and cluster the single-cell transcriptomic atlases of these three species, we identified 13,179 one-to-one gene orthologs between *D. melanogaster* and *D. simulans*. These genes were used to establish anchors for reciprocal principal component analysis (RPCA)-based integration across datasets from all three species (Fig. 1c, Methods). Upon integrating the single-cell transcriptomes of the three species into a unified dataset, we grouped cells into 38 clusters using a shared nearest neighbor (SNN) modularity optimization-based clustering algorithm (Supplementary Fig. 1a,b); notably, this unsupervised clustering did not yield any species-specific cluster, suggesting the global cellular composition of central brains is well-conserved in these species. Our dataset reproduced the previously described global structure of a brain single-cell transcriptome atlas in *D. melanogaster*^21^, where cells are differentiated by their expression of developmental patterning genes (notably, *pros* and *Imp*) (Supplementary Fig. 1c,d). Glial cells (*repo*+) form several clusters, while neurotransmitter markers could distinguish GABAergic (*Gad1*+), monoaminergic (*Vmat*+), cholinergic (*VaChT*+) and glutamatergic (*VGlut*+) neurons (Supplementary Fig. 1e,f).

Within these broad categories of cells, we identified marker genes for each cluster to enable the annotation of central brain cell types. Larger clusters (>∼1,000 cells) were subclustered and marker genes were identified for each subcluster. For example, a monoaminergic neuron cluster was divided into dopaminergic (DOP), serotonergic (SER), and octopaminergic/tyraminergic (OCTY, TY, Tbh) subclusters, and a peptidergic neuron population was divided into subclusters expressing specific neuropeptide genes including *Hug*, *insulin-like peptides*, *Tk*, *Mip* and *Crz* (Supplementary Fig. 2). Using additional marker genes from *D. melanogaster* brain atlases^21,22^ and the expression patterns of genes encoding proteins required for neurotransmission or neuromodulation^42^, we could define 48 annotated and 16 unannotated cell types common to all three species (Fig. 1d,e, Supplementary Fig. 3, Supplementary Table 2, Methods), which together encompassed all cells in the atlases. In addition to previously-characterized cell types^21,22^, the larger size of our combined datasets provided power to identify novel, rare cell types. For example, we identified a cluster expressing the RYamide (RYa) neuropeptide, which comprises only 0.11% of central brain cells (Fig. 1d). Importantly, among the 64 annotated and unannotated cell types, we did not identify any examples of cell types that are unique to any species, or absent in only one species. Our single-cell transcriptomic atlases show that the cellular composition of the central brain is globally conserved across *D. melanogaster*, *D. simulans* and *D. sechellia*, consistent with their overall similarity in gross neuroanatomy (Fig. 1a)^43^.

### Conserved gene expression patterns in drosophilid central brains

We first exploited our atlases to identify genes with conserved expression patterns - beyond the markers used in cell type annotation - which we reasoned would reveal the core molecules with essential roles in global and local brain organization and cellular functions. To identify such genes, we performed correlation analyses of cell type-specific gene expression levels across three pairwise species comparisons: *D. melanogaster*-*D. simulans* (*Dmel-Dsim*), *D. melanogaster*-*D. sechellia* (*Dmel-Dsec*) and *D. simulans*-*D. sechellia* (*Dsim-Dsec*). We focused on the 1,686 genes that are expressed in at least 5% of D. melanogaster central brain cells. The majority of these genes are broadly expressed - in 7 to all 64 cell types (on average, ∼54) - where expression is counted if a gene is detected in at least 5% of the cells of a given type. For each of these genes, we computed the average expression level across 64 cell types and compared these cell type-wide expression patterns across species, subsequently measuring the pairwise correlations (Supplementary Fig. 4a). This analysis identified 368 genes displaying strongly correlated expression patterns (Spearman’s *ρ* > 0.7) across all three species pairings (Supplementary Fig. 4b, Supplementary Table 3). Gene Ontology (GO) term analysis revealed a notable enrichment in genes coding for membrane proteins, including adhesion molecules, ion channels and G-protein coupled receptors (GPCRs), which are likely to play pivotal roles in defining and/or maintaining the structure and function of the nervous system (Supplementary Fig. 4c).

Next, we refined our analysis to genes that are more specifically expressed, positing that these are likely to govern cell type identity or underlie cell type-specific functions. Within each cell type, we identified genes that are consistently expressed in over 30% of cells across all three species. Of these, we focused on the genes that are restricted to only 1-9 of the 48 annotated cell types (an arbitrary cut-off of gene expression specificity; this analysis also excluded the 16 unannotated cell types). With this approach, we cataloged 896 genes exhibiting both conserved and specific expression patterns (Supplementary Table 4). This curated list of genes serves as a valuable resource for linking the functional roles of distinct brain cell types to their unique, yet conserved, gene expression profiles.

The strong conservation of gene expression patterns across species implies the existence of shared gene regulatory mechanisms. While our profiling of mature adult brains is likely to limit our ability to identify important developmental genes, we reasoned that our datasets should still capture information on the expression patterns of terminal selector transcription factors, which establish and maintain the identity of post-mitotic neurons^44,45^. Of the 896 genes displaying specific and conserved expression patterns as described above, 79 encode known or predicted transcription factors. This group includes key developmental regulators, such as a POU domain transcription factor (*acj6*) and a paired-like homeobox transcription factor (*ey*), essential for the development of OPN and Kenyon cells, respectively (Supplementary Fig. 5)^46–48^. This suggests the potential of other genes in this set to play critical roles in regulating cell type identity. For example, another paired-like homeobox transcription factor (*Ptx1*), which is expressed in a few cell types including glutamatergic Fru+ cells (Supplementary Fig. 5), might be a terminal selector for these cell types, similar to its known role for enteroendocrine cell specification in the gut^49^ Furthermore, expression patterns of these transcription factors group cell types with shared functions and/or developmental origins (e.g., glia, Kenyon cells) (Supplementary Fig. 5). Together, these analyses not only define a unique molecular fingerprint for each cell type’s terminal identity but also offer a resource for identifying previously unknown terminal selectors specific to each cell type (Supplementary Table 4).

### Divergence in the frequencies of homologous cell types

Having determined conserved types and molecular properties of likely homologous cell populations in the drosophilid trio, we next explored if and how these species’ brains have diverged. Taking advantage of the statistical power afforded by having six biological replicates per species, we first analyzed interspecific variation in the representation of the 48 annotated and 16 unannotated cell types (Fig. 2a,b, Supplementary Fig. 6). The majority of cell types displayed similar frequencies across the three species, including Kenyon cells, Clock cells and Fruitless cells (which play critical roles in learning and memory, circadian rhythms and sexual behaviors, respectively) (Fig. 2b). Between *D. melanogaster* and *D. simulans*, we did not observe any significant differences in cell type frequencies. However, *D. sechellia* exhibited significant interspecific variation in three cell types: there are ∼3-fold fewer perineurial glial cells (PRN), which comprise the blood-brain barrier (BBB), in this species compared to *D. melanogaster* or *D. simulans* (Fig. 2b,c). By contrast, this specialist species has higher frequencies of the excitatory short neuropeptide F-producing cell type (sNPF(E)) compared to the other species (Fig. 2a-e). sNPF(E) cells encompass diverse subtypes (Supplementary Fig. 7) and some of these exhibit higher frequencies in *D. sechellia*, while others have conserved representation across the three species (Fig. 2c-e). Finally, we also observed a higher proportion of Dh44 neuroendocrine cells in *D. sechellia* compared to *D. melanogaster* (Fig. 2a-c). However, an increased, albeit non-significant, frequency of Dh44 neurons was also noted in *D. simulans* (Fig. 2a-c), suggesting that expansion in this cell population took place before speciation of *D. sechellia* and/or that there was a reduction in this population in the *D. melanogaster* lineage. Together our results suggest that the cellular composition of the central brain in *D. sechellia* is highly conserved with its generalist relatives, but a few cell types might have changed during ecological specialization.

**Figure 2.**
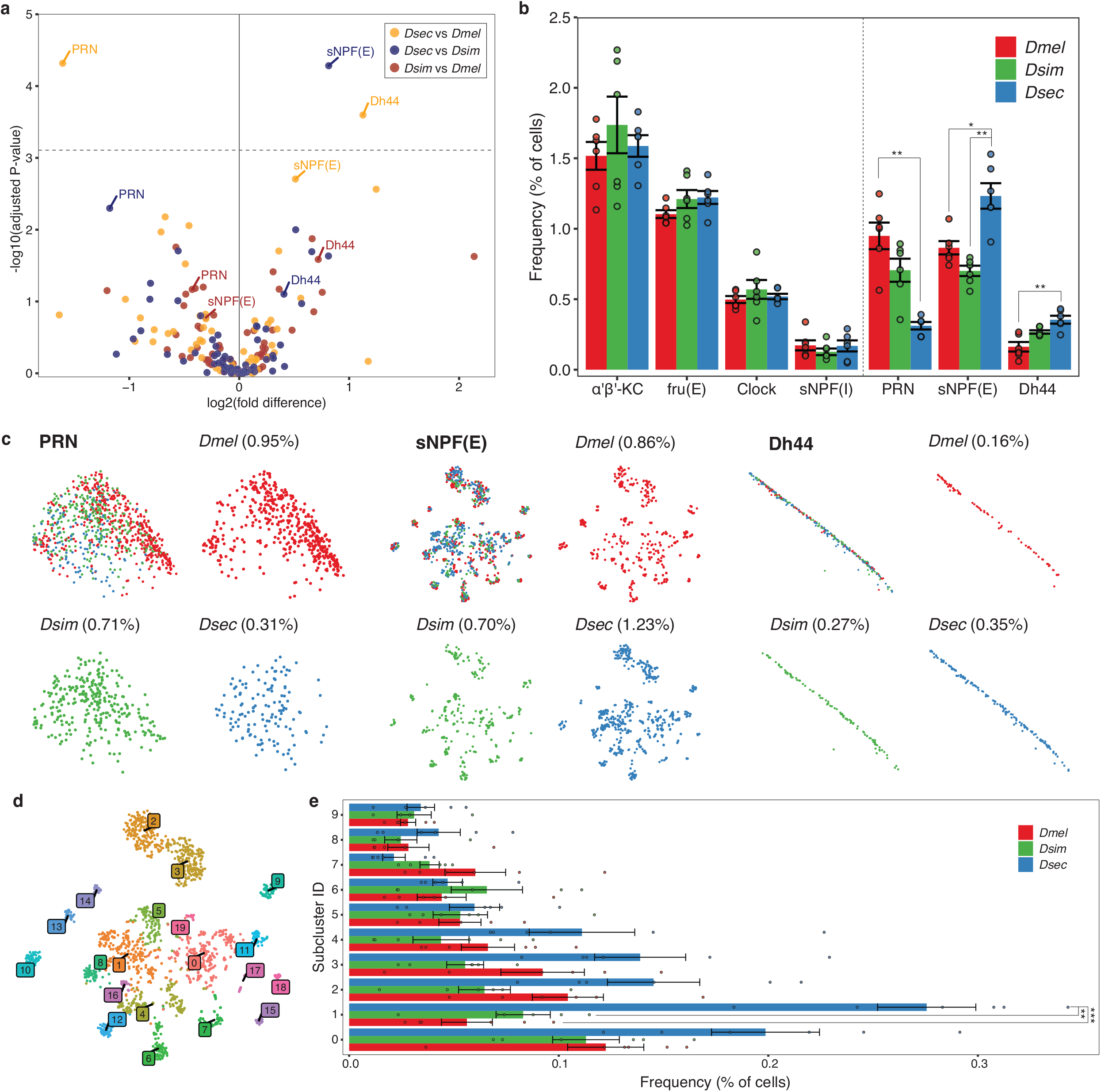
Species divergence in the frequencies of central brain cell types. (a) Volcano plot of the pairwise comparisons of cell type frequencies in the central brains of the three drosophilid species. The dashed horizontal line denotes the Bonferroni-corrected *P*-value threshold. (b) Bar plot illustrating selected cell type frequency comparisons. To the left of the dashed vertical line are four cell types with no significant frequency differences across the three species. To the right, three cell types exhibit significant frequency variations in at least one pairwise species comparison. Each point corresponds to one of the six biological replicates. (c) tSNE plots of PRN, sNPF(E) and Dh44 populations. Cells are colored by their species of origin. The frequency of each cell type within its respective species is indicated in parentheses. The sNPF(E) plots were produced from the dataset after subclustering of sNPF(E) cells. (d) tSNE plot of the sNPF(E) cells colored by the SNN-based clustering. (e) Bar plots showing comparisons of cell type frequencies of the top ten largest subclusters of sNPF(E) cell populations. Each point corresponds to one of six biological replicates. Statistical significance in (b,e) was calculated by One-Way ANOVA with Tukey’s HSD post-hoc test. *P*-values were adjusted using false discovery rate (FDR) for multiple comparisons. \*\*\**P* < 0.001; \*\**P* < 0.01; * *P* < 0.05.

### Transcriptomic divergence in homologous cell types across the drosophilid trio

Given the global conservation of cell type and frequency in these drosophilids’ brains, we reasoned that phenotypic differences between these species might be reflected more prominently in the divergence of the transcriptomic profile of homologous cell types. We identified the 50 most abundantly expressed genes from each of the 48 annotated cell types in *D. melanogaster* and examined the transcriptomic similarity between species for homologous cell types using correlation analysis (Fig. 3a-c). The similarity of gene expression profiles varies across cell types. For example, gene expression levels are highly similar in insulin-producing cells (IPC) (*Dmel-Dsim*: *ρ* = 0.76, *Dmel-Dsec*: *ρ* = 0.74, *Dsim-Dsec*: *ρ* = 0.89) compared to PRN (*Dmel-Dsim*: *ρ* = 0.53, *Dmel-Dsec*: *ρ* = 0.39, *Dsim-Dsec*: *ρ* = 0.74). Globally, cell-type transcriptomic divergence reflects the phylogenetic distance of the different pairs: the more closely-related *Dsim-Dsec* pair shows higher similarity than both the *Dmel-Dsim* and *Dmel-Dsec* pairs across all 48 cell types (Fig. 3c). Although most of the cell types show a similar degree of transcriptomic divergence between *Dmel-Dsim* and *Dmel-Dsec* pairs, we noted that a few cell types display substantial differences between two pairs (Fig. 3c). Notably, four out of five glial cell types (astrocytes (AST), cortex glia (CTX), PRN and subperineurial glia (SUB)) show reduced similarity in the *Dmel-Dsec* pair than the *Dmel-Dsim* pair (Fig. 3b,c), suggesting that gene expression profiles of these glial cell types have changed during host specialization along the *D. sechellia* lineage.

**Figure 3.**
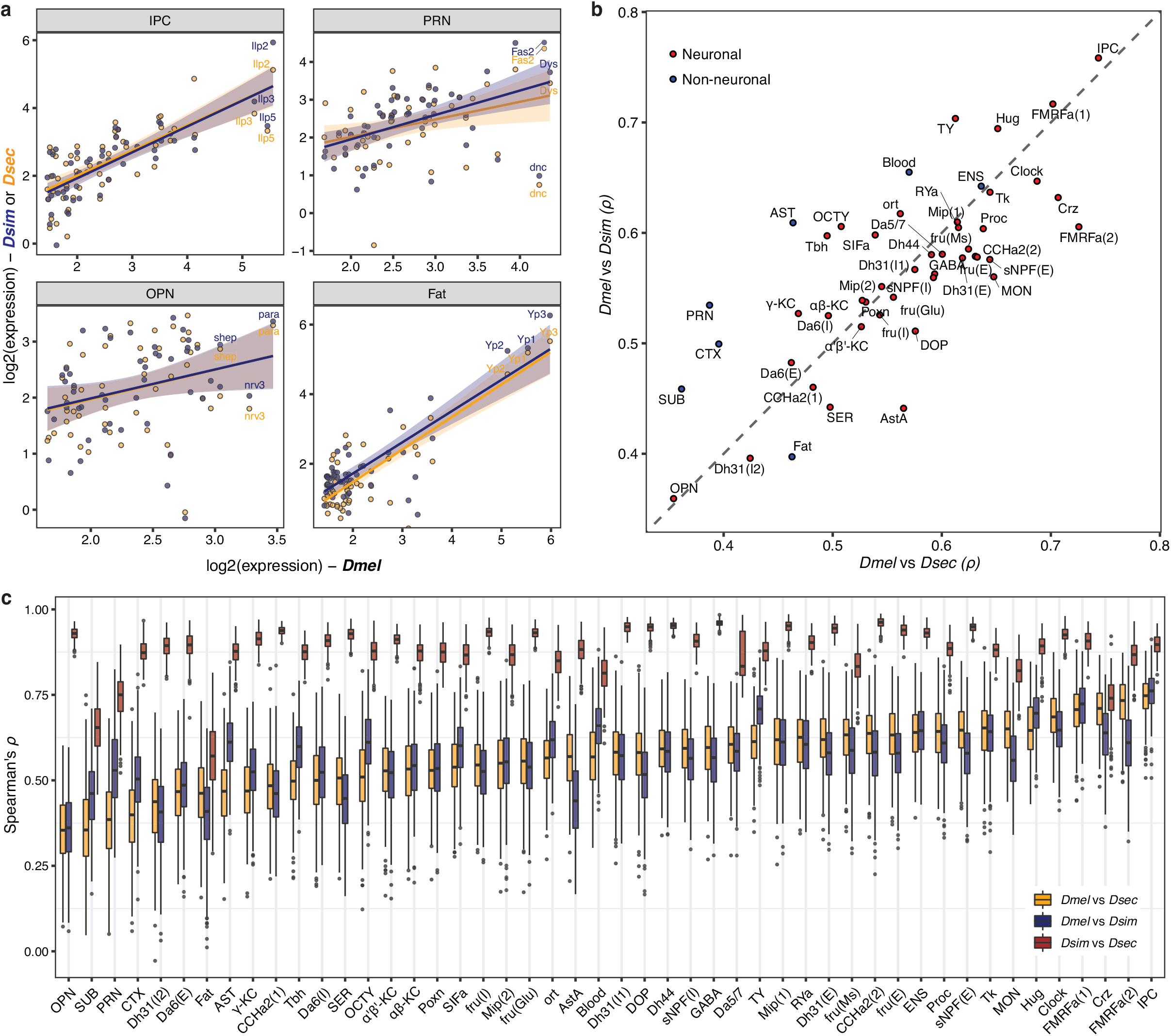
Gene expression divergence of conserved central brain cell types across three drosophilid species. (a) Scatter plots illustrating the interspecific gene expression variations across four distinct cell types: IPC (insulin-producing cells), PRN (perineurial glia), OPN (olfactory projection neurons) and Fat (fat body, an adipocyte-like tissue). The analysis features the 50 genes (each represented by a dot) with the highest expression levels in the pseudobulk transcriptome of *D. melanogaster* for each cell type. The plots display expression levels of these genes for three drosophilid species: *D. melanogaster* on the x-axis, and *D. simulans* (navy) and *D. sechellia* (orange) on the y-axis. Smoothed lines are linear model fits. (b) Scatter plot illustrating the transcriptomic similarities among 48 annotated cell types between *D. melanogaster* and *D. sechellia* (x-axis), and *D. melanogaster* and *D. simulans* (y-axis). Each point represents an individual cell type, with neuronal cell types highlighted in red and non-neuronal ones in blue. Spearman correlation coefficients (*ρ*) are derived from the expression level similarities of the top 50 highly-expressed genes in the pseudobulk transcriptome of *D. melanogaster* for each respective cell type. (c) Tukey box plots presenting transcriptomic similarities among 48 annotated cell types in pairwise comparisons of three drosophilid species. The Spearman correlation coefficients (*ρ*) are calculated based on the expression similarities of 30 randomly selected genes (from 400 permutations) among the top 50 highly-expressed genes in *D. melanogaster*’s pseudobulk transcriptome for each cell type. The horizontal line in the middle of each box is the median; box edges denote the 25th and 75th quantiles of the data; and whiskers represent 1.5× the interquartile range.

To examine transcriptomic divergence at higher resolution, we next characterized differentially expressed genes (DEGs, fold-change threshold >1.5, adjusted P-value <0.05) within each cell type across the three species (*Dmel-Dsec*: n=487, *Dmel-Dsim*: n=369, *Dsim-Dsec*: n=242) (Supplementary Table 5). Different cell types exhibited different DEGs, as illustrated for PRN and sNPF(E) cells (Fig. 4a,b). Importantly, the vast majority of DEGs were identified as divergently expressed from only 1-3 cell types (Fig. 4c). For example, among 487 DEGs from the *Dmel-Dsec* pair, 275 genes (56.5%) were identified from a single cell type, and 421 genes (86.4%) were identified from no more than 3 cell types, even though the majority of these genes are broadly expressed (on average, expressed in 45/64 cell types in *D. melanogaster*, minimum percent expression threshold: 5%). We next examined the cell type identity for each DEG (Fig. 4d). Notably, in all species comparisons, four glial cell types (CTX, PRN, AST and ensheathing glia (ENS)) consistently exhibited the highest number of DEGs, suggesting that, in comparison to their neuronal counterparts, glial cell types tend to display more divergent gene expression profiles across species. Specifically, cortex glia (CTX), which ensheath neuronal cell bodies in the brain cortex and are closely associated with the BBB^50^, show the largest number of DEGs for all three comparisons. Among the annotated neuronal cell types, those expressing the neuropeptide Allostatin A (AstA) displayed the greatest number of DEGs in two comparisons (*Dmel-Dsec*, *Dsim-Dsec*) and the second highest in the third (*Dmel-Dsim*). The observed cross-cell type and cross-species variation in transcriptomic divergence suggests that each cell type has undergone unique changes during the evolution of these species.

**Figure 4.**
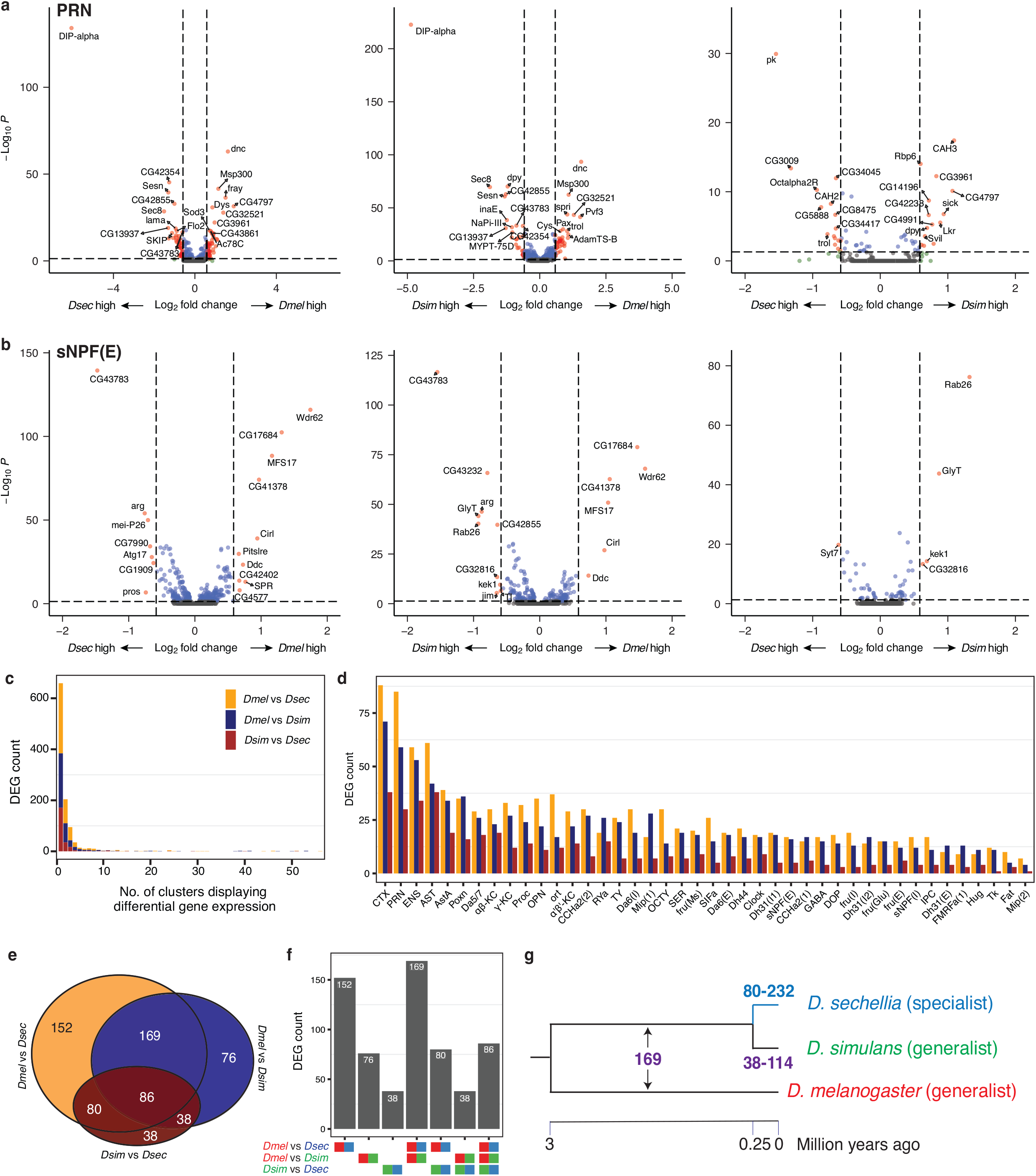
Characterization of differentially expressed genes (DEGs) in conserved cell types across *Drosophila* central brains. (a,b) Volcano plots showing DEGs (red) in two specific cell types, PRN (a) and sNPF(E) (b), across three pairwise species comparisons. For comparisons with more than 20 DEGs, only the top 20 most significant genes are labeled. (c) Histogram displaying the frequency of DEGs sorted by the total number of cell types/clusters in which they are identified as DEGs. Stacked bars with distinctive colors represent DEGs identified in pairwise species comparisons. (d) A bar plot (color-coded as in (c)) showing the frequency of DEGs across 42 cell types with at least one DEG identified from all pairwise comparisons among three species. Cell types are arranged in order of the number of DEGs across three pairwise comparisons. (e,f) A Venn diagram (e) (color coded as in (c)) and a bar plot (f) illustrating the intersections of DEGs identified in three pairwise species comparisons. (g) Hypothetical numbers of genes that underwent expression changes are shown across the lineages of *D. melanogaster*, *D. simulans* and *D. sechellia*.

### Unique gene expression changes in *D. sechellia*

To investigate gene expression changes potentially related to the ecological specialization of *D. sechellia*, we analyzed the overlap of DEGs from different species comparisons. Overall, we found that 58% of DEGs were identified in more than one comparison (Fig. 4e,f). For example, a set of 169 DEGs appeared in both *Dmel-Dsec* and *Dmel-Dsim*, but not in *Dsim-Dsec*, suggesting these shared DEGs may reflect expression changes originating either in the *D. melanogaster* lineage or in the common ancestor of *D. simulans* and *D. sechellia* (Fig. 4g). Importantly, we observed 32% more DEGs from *Dmel-Dsec* (n=487) than *Dmel-Dsim* (n=369), indicating that the *D. sechellia* lineage has gained more gene expression changes (Fig. 4e). When we inferred lineage-specific gene expression changes by investigating overlaps among DEGs (Fig. 4e,f, Methods), we found that the estimated number of *D. sechellia* DEGs (n=80-232) far exceeded *D. simulans* DEGs (n=38-114) (Fig. 4e-g). Given that a similar number of DEGs would be expected if expression differences were solely due to divergence time (’neutral changes’), this observation suggests that most DEGs in the *Dsim-Dsec* pair likely result from expression changes specific to the *D. sechellia* lineage.

We next focused on the 80 genes with expression changes specific to the *D. sechellia* lineage (Supplementary Table 6). As the number of DEGs was too limited for traditional GO term enrichment analysis, we manually curated their putative/known functions. In neuronal cell types, we identified 39 *D. sechellia* DEGs including those encoding aminergic GPCRs such as a serotonergic receptor (*5-HT1A*) and two octopaminergic receptors (*Octbeta2R*, *Octbeta3R*), a prohormone convertase (*amon*), a cyclic nucleotide phosphodiesterase (*Pde1c*), and a glucose transporter (*Glut1*). These neuronal *D. sechellia* DEGs are identified from a large fraction of the cell types, where AstA cells and α’β’ Kenyon cells show the largest number of DEGs (n=5) (Fig. 5a).

**Figure 5.**
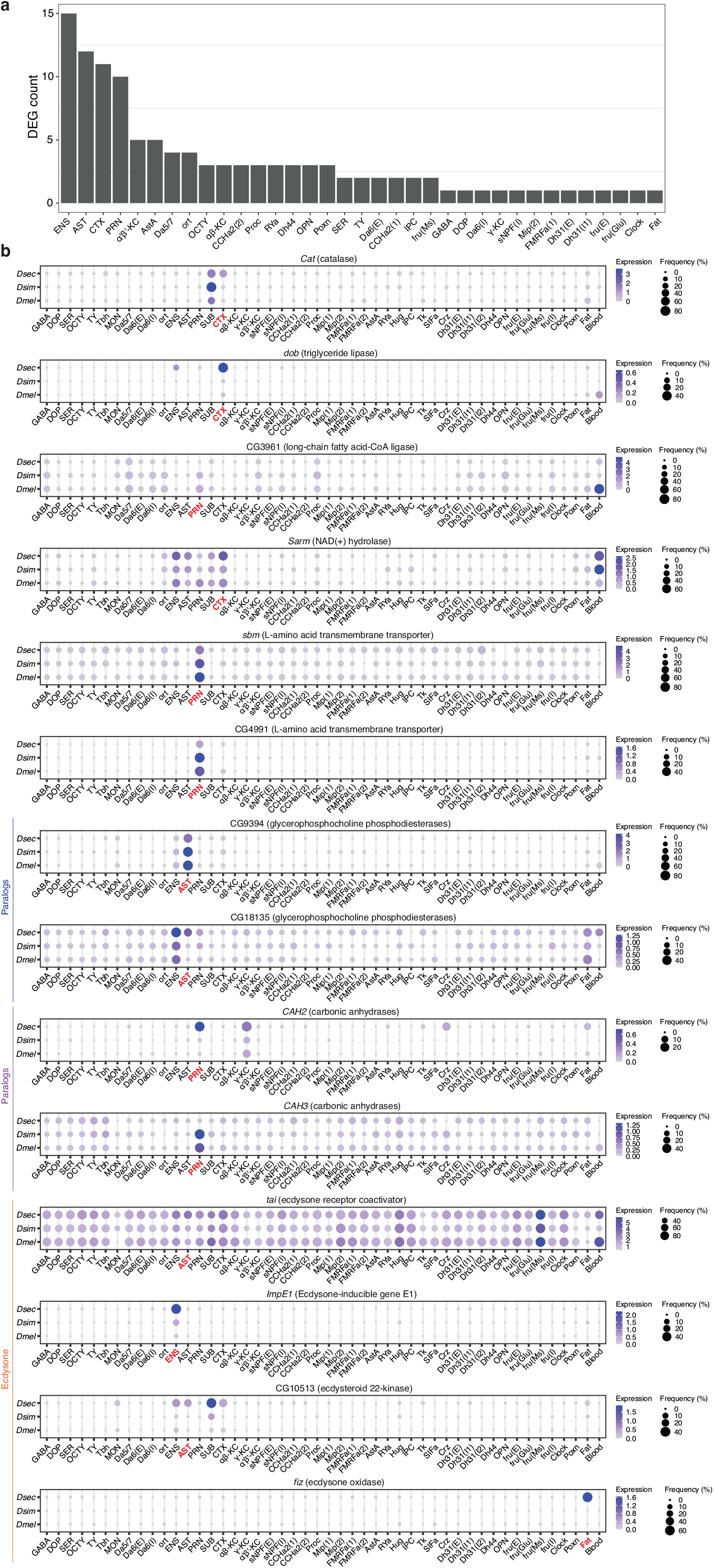
Divergent gene expression in *D. sechellia*. (a) Histogram of the frequency of DEGs across 48 cell types from *D. sechellia*-specific DEGs. Cell types are arranged by decreasing DEG number. (b) Dot plots illustrating expression levels and frequencies of 14 *Dsec*-specific DEGs in *D. melanogaster*, *D. simulans* and *D. sechellia* across 48 cell types. Cell types are shown on the x-axis and species are shown on the y-axis.

Among non-neuronal cell types, we identified 40 *D. sechellia* DEGs in four glial cell types (ENS, AST, PRN, CTX) (Fig. 5a), consistent with the pronounced interspecific gene expression variation in glia (Fig. 4d). These glial DEGs are enriched with various metabolic genes (Fig. 5b); including catalase (*Cat*), triglyceride lipase (*dob*), long-chain fatty acid-CoA ligase (CG3961), NAD(+) hydrolase (*sarm*), L-amino acid transmembrane transporter (*sbm* and CG4991), glycerophosphocholine phosphodiesterases (CG9394 and CG18135) and carbonic anhydrases (*CAH2* and *CAH3*). Intriguingly, for both of the latter two paralog pairs we noted opposing expression changes in *D. sechellia*: CG9394 is downregulated but CG18135 is upregulated in AST, and *CAH2* is upregulated and *CAH3* downregulated in PRN. These observations imply potential gene expression dosage compensation between paralogs.

Beyond metabolic genes, we identified three glial *D. sechellia* DEGs involved in ecdysone signaling: ecdysone receptor coactivator (*tai*), Ecdysone-inducible gene E1 (*ImpE1*) and ecdysteroid 22-kinase (CG10513) (Fig. 5b). Furthermore, the ecdysone oxidase gene, *fiz*^51^, displayed specific expression in the fat body (Fat) tissue of *D. sechellia*, while absent in both *D. melanogaster* and *D. simulans* central brains (Fig. 5b). This gene had previously been highlighted as one of the most divergently expressed genes between the heads of *D. simulans* and *D. sechellia*^52^, and experimental evolution of *D. melanogaster* populations under a novel dietary condition reproducibly resulted in changes to the expression of *fiz* ^51^. These findings collectively imply that ecdysone signaling might have played a pivotal role in the genetic adaptation to distinct dietary conditions in the *D. sechellia* lineage.

### Gene expression plasticity in the specialist brain

While these comparative atlases were intentionally generated from flies grown on the same food medium, we considered the possibility that the composition and transcriptome of the *D. sechellia* brain might be influenced by the presence of nutrients in noni. Supplementing food with noni juice paste greatly improved this species’ fitness under laboratory conditions, as assessed by egg number and development to the pupal stage (Fig. 6a). In parallel with the datasets described above, we also generated a central brain cell atlas from *D. sechellia* that had been reared on standard medium supplemented with noni paste. Comparisons of the snRNA-seq central brain transcriptomes of *D. sechellia* grown with or without noni supplement revealed essentially no effect on the cellular composition of the *D. sechellia* central brain, except a small increase in the frequency of Clock neurons (Supplementary Fig. 8). Moreover, examination of gene expression differences using the same threshold that yielded hundreds of interspecific DEGs, we identified only one gene, that is more highly expressed in the brains of flies grown on noni: CG5151, which encodes a Low-Density Lipoprotein Receptor Class A Domain Containing 4 (LDLRAD4) homolog (Fig. 6b,c). Interestingly, CG5151 is broadly-expressed but differentially expressed specifically in PRN (Supplementary Fig. 9), which is one of the cell types with the most significantly changed frequency and gene expression between species (Fig. 2a-c and Fig. 4d).

**Figure 6.**
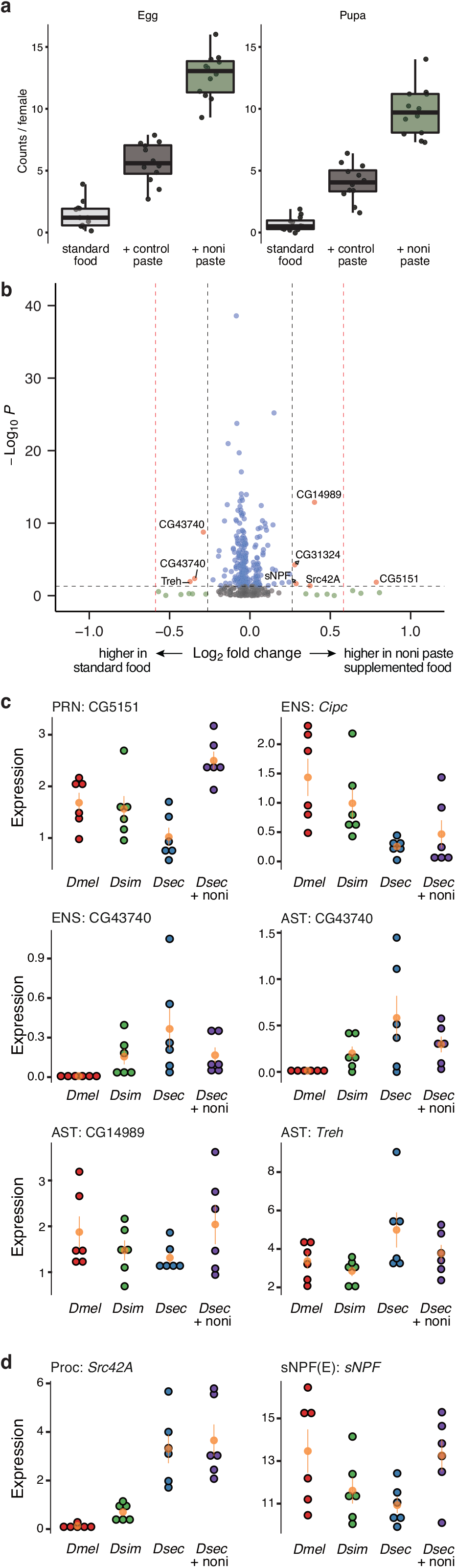
*D. sechellia* brain gene expression changes due to noni. (a) Tukey box plots for brood size (left) and pupal count (right) per mated *D. sechellia* female grown in three conditions. Each point corresponds to a biological replicate. The horizontal line in the middle of each box is the median; box edges denote the 25^th^ and 75^th^ quantiles of the data; and whiskers represent 1.5× the interquartile range. (b) Volcano plot highlighting differences in gene expression of *D. sechellia* central brains under two distinct growth conditions: with and without added noni paste. All cell type-specific DEG analyses are integrated into this single representation, and a reduced fold-change threshold (from 50% to 20%) is applied for the identification of DEGs. (c-d) Dot plots illustrating expression levels of eight plasticity genes in glial (c) and neurons (d) in response to noni treatment. The x-axis displays species and growth conditions, while expression levels are represented on the y-axis.

When we lowered the fold-change threshold (to >1.2), we identified 6 more noni DEGs (Fig. 6b). Four of these (CG14989, CG43740, *Cipc* and *Treh*) (Fig. 6c) are differentially expressed in glial cell types, suggesting that glia are not only diverged in their gene expression between *D. sechellia* and two generalist *Drosophila* species, but are also the most responsive to environmental conditions that drove the evolution of *D. sechellia* lineage. For example, expression of a trehalase (*Treh*) gene is higher in AST glia in *D. sechellia* than *D. melanogaster* or *D. simulans* when grown in the same standard medium; but *Treh* expression is decreased when noni juice is supplemented to *D. sechellia*. The other two noni DEGs are expressed in neurons: *Src42A* and *sNPF*, from Proc and sNPF(E) cell types, respectively (Fig. 6d). Among the three species grown in the standard medium, *D. sechellia* showed the lowest *sNPF* expression in sNPF(E) cells; but it recovered its expression level comparable to generalist species upon noni juice supplementation.

## Discussion

Our work provides an unprecedented comprehensive view of how animal brains evolve in a well-defined phylogenetic and ecological framework, by taking advantage of the relatively small brains of closely-related drosophilid species to generate and compare whole central brain cellular atlases. Previous studies on how the nervous system of *D. sechellia* differs from *D. melanogaster* and *D. simulans* have primarily focused on peripheral chemosensory pathways^28,32–34,36–38,40^, leaving knowledge of potential adaptations in the brain almost completely unexplored. Despite the very different ecology and behaviors of the equatorial island-endemic specialist *D. sechellia* from its cosmopolitan, generalist relatives, the overall brain architecture of these flies is highly conserved. Within the resolution of our clustering analyses, we did not detect any species-specific cell population. Moreover, the frequencies of the vast majority of presumed homologous cell types are conserved. This observation contrasts with studies of the peripheral olfactory system, where several olfactory sensory neuron population are increased (or decreased) in *D. sechellia* compared to the other species^32,33,38,40^. Our data provide empirical data to help answer the long-held question as to whether the sensory periphery is more evolvable than central brain regions^5^. The high conservation of central brain cell populations might reflect their more pleiotropic functions compared to individual sensory neuron populations.

In light of this conservation, consideration of the central cell populations that do differ between species is of interest. The most pronounced difference in *D. sechellia*’s central brain composition was found not in neurons but in a specific glial type, PRN, which forms a diffusion barrier around the nervous system as part of the BBB. In *D. melanogaster*, PRNs have an important role in sugar uptake into the brain^53^. Notably, noni fruit has a much lower carbohydrate (sugar):protein ratio than most other fruits, and *D. sechellia* cannot develop on carbohydrate-rich diets, apparently due to defects in carbohydrate-induced metabolic and gene expression changes^54^. We speculate that the reduction in PRNs in this species reflects a reduced requirement for sugar uptake by the brain. While it is unclear whether this loss is adaptive in *D. sechellia* (i.e., increasing fitness of this species in another, unknown, way), it might be a phenotype that constrains this species to this niche (i.e., representing an “evolutionary dead end”^55^). Of the neurons, only the excitatory sNPF-expressing population is significantly different in *D. sechellia* compared to both generalist species. *D. melanogaster* sNPF is expressed in numerous groups of neurons located in various regions of the brain^56^, commensurate with the many subclusters of sNPF(E) we identified in our atlases. This neuropeptide has been linked to regulation of diverse behaviors, including olfaction, feeding, circadian rhythms and sleep, as well as organismal growth and lifespan^57,58^. In most of these roles, the neuronal population(s) in which sNPF is required has not, however, been delineated. Dissection of the contribution, if any, of subsets of expanded sNPF(E) neuron subtypes in *D. sechellia* to species-specific behaviors will require the means to selectively visualize and manipulate specific cell types in both this species and *D. melanogaster*.

Within homologous cell types, global patterns of gene expression – spanning those encoding known (or newly predicted) terminal transcription factors to signaling effectors, such as neurotransmitters/peptides – are, as expected, conserved. However, we observed many examples of species-specific divergence in gene expression, which was different in each cell type in terms of gene identity and magnitude of change. In particular, we found that many broadly expressed genes differ in their expression between species only in a subset of specific cell types. Such properties reinforce the idea of cell types being independent evolutionary units^1^. Of particular interest is our observation that the majority of glial cell types display the greatest expression divergence. These properties might reflect less stringent selective pressures on glia to maintain precise structure and function compared to neurons that act within stringently-defined circuitry. However, the enrichment of gene expression changes in glia might also reflect adaptive changes within these cell types, which have numerous known or presumed supportive roles in the brain, such as in homeostatic regulation and filtering of the external milieu for energy supply to neurons and waste removal^59^.

It is important not to interpret every change in gene expression as indicative of adaptive evolution: after species divergence, neutral genetic polymorphisms accumulate, raising the possibility that gene expression also undergoes neutral evolution^60,61^. Our analysis of the drosophilid trio offers an ideal model system to distinguish neutral and potentially non-neutral gene expression evolution. The expectation, under neutral conditions, is for *D. simulans* and *D. sechellia* to exhibit a similar degree of gene expression divergence when compared to *D. melanogaster*. However, our data revealed more pronounced shifts in brain gene expression in *D. sechellia*, suggesting this lineage underwent non-neutral gene expression changes, possibly due to its adaptation to its unique ecological niche. Glia stand out as the cell types displaying the most potential adaptive expression alterations, reinforcing the notion that they have an important contribution to brain evolution. Among the genes displaying altered expression patterns, those associated with metabolism and the ecdysone signaling pathway are particularly enriched. The metabolic gene regulatory network likely underwent remodeling to adapt to the new nutritional conditions of the species’ unique niche. Ecdysone signaling coordinates nutritional status with organ growth and patterning in *D. melanogaster*^62^; it is possible that changes in hormonal signaling occurred in the *D. sechellia* lineage to allow alignment of developmental regulation with its novel nutritional conditions. Supporting the hypothetical adaptive nature of interspecific variation in ecdysone signaling, a previous study highlighted that ecdysone-regulated genes with variable expression across species tend to be under lineage-specific selection, as opposed to undergoing neutral evolution^60^.

One challenge of comparing transcriptomes of ecologically-diverse species is distinguishing between genetic and environmental effects on gene expression. To minimize environmental influences, our initial comparative atlases were of the drosophilid species grown on an identical food medium. However, as *D. sechellia* is thought to have metabolic defects^31,54^, we also generated a brain atlas from flies grown with noni juice. Although this medium substantially increased *D. sechellia*’s survival and fecundity, there was very little influence upon the cellular composition or gene expression patterns in the brain of this species. These results support the idea that the interspecific differences observed are largely genetically-determined. Of the observed environmental-dependent gene expression changes, the majority occur in various types of glia, emphasizing the plasticity of these cell types both over evolutionary timescales and in response to environmental changes. It is most noteworthy that the glial gene (CG5151) and neuronal gene (*sNPF*) displaying the largest differential expression in *D. sechellia* cultured with and without noni are markers of the two main cell types (PRN and sNPF(E), respectively) that display differential population size between species. These observations hint at the possibility that the cellular nodes in the brain that initially responded to environmental presence of noni in the *D. sechellia* ancestor eventually have been reshaped by natural selection during species-specific niche adaptation^63^.

## Methods

### *Drosophila* strains and culture

Flies used in this study were *D. melanogaster* (Canton-S), *D. simulans* (*Drosophila* Species Stock Center (DSSC) 14021-0251.004) and *D. sechellia* (DSSC 14021-0248.07), which were all cultured on standard wheat flour–yeast–fruit juice food. For the animals analyzed in Figure 6, *D. sechellia* was also grown on standard food supplemented on top with a paste of Formula 4-24 Instant *Drosophila* medium (blue, Carolina Biological Supply) mixed 1:5 weight:volume in noni juice (Raab Vitalfood).

### Single-nucleus RNA sequencing (snRNA-seq)

30-50 newly-eclosed male and female adult flies were collected and placed together for mating, followed by sorting by sex on day 5. The central brains of 20 mated female flies were dissected and collected in 100 μl Schneider’s medium, flash-frozen in liquid nitrogen, and stored at −80°C for nuclear extraction. Sample homogenization, single-nucleus suspension, and nuclear sorting were performed using the standard protocol described in the Fly Cell Atlas^23^. To obtain sequencing data from 10,000 nuclei, 15-20,000 nuclei were collected and loaded onto the Chromium Next GEM Chip (10x Genomics). Sequencing libraries were prepared with the Chromium Single Cell 3ʹ reagent kit v3.1 dual index, strictly following the manufacturer’s recommendations. Libraries were quantified by a fluorimetric method and their quality was assessed on a Fragment Analyzer (Agilent Technologies). Cluster generation was performed with 0.8-1.0 nM of an equimolar pool from the resulting libraries using the Illumina HiSeq 3000/4000 PE Cluster Kit reagents. Sequencing was performed on the Illumina HiSeq 4000 using HiSeq 3000/4000 SBS Kit reagents according to 10x Genomicsʹ recommendations (28 cycles read1, 8 cycles i7 index read, 8 cycles i5 index, and 91 cycles read2). Sequencing data were demultiplexed using the bcl2fastq2 Conversion Software (v2.20, Illumina). Six biological replicates were processed resulting in snRNA-seq data from 120 central brains each for *D. melanogaster*, *D. simulans* and *D. sechellia* grown on standard food, and *D. sechellia* grown on noni-supplemented food.

### Generation and integration of single-cell central brain transcriptomic atlases

Raw snRNA-seq data was first processed through Cell Ranger (v6.0.1, default parameters except--include-introns)^41^. *D. melanogaster* reference genome and transcriptome from FlyBase (release 6.34) were used for all three species (*Dmel*-to-*Dmel*, *Dsim*-to-*Dmel*, *Dsec*-to-*Dmel*); additionally, sequence reads from *D. simulans* and *D. sechellia* were also processed with the *D. simulans* reference genome and transcriptome from FlyBase (release 2.02) (*Dsim*-to-*Dsim, Dsec*-to-*Dsim*). Gene expression matrices from *Dsim*-to-*Dmel* and *Dsec*-to-*Dmel* processing were used for differential gene expression analysis (see below). All gene expression matrices were processed with SoupX (v1.6.1, default parameters) ^64^ to remove ambient RNA contamination. Decontaminated *Dmel*-to-*Dmel*, *Dsim*-to-*Dsim*, and *Dsec-*to-*Dsim* datasets were then normalized through SCT normalization^65^ and putative doublets were filtered out using DoubletFinder (v2.0.3)^66^. Decontaminated and filtered datasets were subsetted for 13,179 one-to-one orthologs between reference genomes of *D. melanogaster* and *D. simulans*, which were inferred using Orthofinder^67^ and manual curation of reciprocal best blastp hits; subsetted datasets were then integrated through the reciprocal PCA (RPCA) based integration method implemented in Seurat (v4.3.0, functions SelectIntegrationFeatures (nfeatures = 3000), PrepSCTIntegration, FindIntegrationAnchors (reference=*D. melanogaster*) and IntegrateData)^68^. PCA was performed on this integrated dataset; the first 50 PCs were used for clustering of cells in the integrated dataset into 38 clusters (functions FindNeighbors and FindClusters, default parameters except for resolution=0.2) and generation of tSNE and UMAP plots.

### Cell type annotation and quantification

Cell clusters were annotated using marker genes obtained through the FindMarkers function in Seurat (only.pos = TRUE, min.pct = 0.15, logfc.threshold = 0.25, test.use = “MAST”). Larger clusters (representing >0.5% of total cells sequenced (i.e., >∼1,000 cells) underwent further subclustering, with marker genes identified for each subcluster. Cluster and subcluster identities were primarily determined by comparing these marker genes with those identified in previous studies for *D. melanogaster*^21,22^. Any clusters and subclusters that remained unannotated following this step were then identified based on marker genes involved in neurotransmission and neuromodulation^42^. This process resulted in the annotation of 22 clusters and 26 subclusters, each identified as distinct cell types. The 16 clusters that remained unannotated after these steps were classified as unannotated cell types. The frequency of each cell type was determined by quantifying the proportion of cells assigned to a specific cell type relative to the total number of cells in each scRNA-seq experiment, with six replicates per species/condition. The significance of interspecific variation in cell type frequency was calculated by a one-way analysis of variance (ANOVA) test, followed by Tukey’s post hoc test for multiple comparisons.

### Conserved gene expression analysis

To mitigate artifacts stemming from the use of different reference genomes, we selected genes that exhibited similar expression levels when aligned to multiple genomes. Specifically, reads from *D. simulans* and *D. sechellia* were aligned to both the *D. melanogaster* and *D. simulans* genomes. Subsequently, the percent expression for every gene was calculated across all cells, and ranks between the two alignments (one to the *D. melanogaster* genome and the other to the *D. simulans* genome) were compared. Of 1,953 genes that exhibited less than a 5% rank difference for both *D. simulans* and *D. sechellia* samples, we retained 1,686 genes that were expressed in at least 5% of *D. melanogaster* central brain cells for the analysis. The AverageExpression function in Seurat was employed to calculate the average gene expression of these selected genes for each cell type. Expression levels of these genes across 64 annotated and unannotated cell types were then compared across species, with the similarity between species being assessed through correlation analysis using Spearman’s *ρ*. FlyEnrichr (https://maayanlab.cloud/FlyEnrichr/) was used for the GO analysis on 368 genes with conserved expression patterns^69,70^.

### Transcriptomic comparisons in homologous cell types

Before comparing gene expression between homologous cell types, we selected genes with reliable expression level information for every cell type by excluding genes whose expression levels in *D. simulans* or *D. sechellia* showed discrepancies (5% rank difference) when their transcripts were aligned to either *D. melanogaster* or *D. simulans* reference genomes. As a result of these criteria, we omitted 890 genes (10.8% of analyzed genes). Subsequently, we identified the top 50 most abundantly expressed genes within each of the 48 annotated cell types in *D. melanogaster*. Transcriptomic similarities between species for homologous cell types were assessed through correlation analysis.

### Differential gene expression analysis

For cell type-specific DEG analysis, cells were subsetted based on their membership and grouped by species and growth conditions (four groups: *Dmel*, *Dsim*, *Dsec*, and *Dsecnoni*). DEGs were identified using the FindMarkers function in Seurat (test.use = “MAST”, min.pct = 0.05). To reduce false discoveries of DEGs that might arise from discrepancies or errors in the genome annotation of different species, sequence reads from *D. simulans* and *D. sechellia* were mapped to the reference genomes of both *D. melanogaster* and *D. simulans*. A gene was only designated as differentially expressed if the mapping of the gene to both genomes consistently indicated differential expression (adjusted *p*-value < 0.05, fold difference > 50%) between the species or condition. For gene expression plasticity analysis, a lower fold-difference threshold (>20%) was also examined.

### Lineage-specific gene expression changes

To infer gene expression changes that occurred after the divergence of *D. simulans* and *D. sechellia* lineages, two methodologies were implemented. The initial approach comprised of identifying non-overlapping DEGs between *Dmel-Dsim* DEGs and *Dmel-Dsec* DEGs, under the assumption that these DEGs do not encompass gene expression differences between the *Dmel* lineage and *Dsim*/*Dsec* lineage before their divergence. This method established the highest counts for *D. simulans* and *D. sechellia* DEGs. The second approach identified the intersection between DEGs of *Dsim-Dsec* and *Dmel-Dsim* or *Dmel-Dsec*. This approach assumed that any *Dsim* or *Dsec* lineage-specific changes would be captured by overlap between *Dsim-Dsec* and *Dmel-Dsim* (*D. simulans* DEGs) or *Dsim-Dsec* and *Dmel-Dsec* (*D. sechellia* DEGs), respectively. This approach established minimum counts for *D. simulans* and *D. sechellia* DEGs.

### Fitness assay

The fitness of *D. sechellia* was measured under three conditions: standard food, standard food supplemented with Formula 4-24 Instant *Drosophila* Medium blue (Carolina Biological Supply) that was hydrated with distilled water at a 1:5 weight-to-volume ratio (“Control paste” in Fig. 6a), or with noni paste (described above; “Noni paste” in Fig. 6a). For each condition, ten mated, 5-day old females were introduced into a culture vial. Brood sizes were measured after 24 h, followed by counting the number of pupae on day 8-9.

## Data availability

Raw sequencing data and processed expression matrices are archived in NCBI Gene Expression Omnibus (GEO) under accession code GSE247965. Processed single-cell transcriptomic atlases are available from SCope (https://scope.aertslab.org/).

## Code availability

All datasets and code for generating the figures and tables are available from GitHub (https://github.com/Evomics/FlyBrainEvo).

## Acknowledgements

We thank the Flow Cytometry Facility (D. Labes) and Lausanne Genomic Technologies Facility (J. Marquis, C. Peter, R. Sermier and K. Bojkowska) of the University of Lausanne for assistance with cell preparation and sequencing. We are grateful to members of the Benton laboratory for discussions and comments on the manuscript. D.L. is supported by the National Research Foundation of Korea (NRF) grants funded by the Ministry of Science and ICT under Project Numbers RS-2023-00211007 and RS-2023-00218602, and by the Ministry of Education under Project Number NRF-2019R1A6A1A10073079. Research in R.B.’s laboratory is supported by the University of Lausanne, an ERC Advanced Grant (833548) and the Swiss National Science Foundation (310030B_185377).

## Author contributions

D.L. and R.B. conceived the project. D.L. generated all datasets and performed all analyses, with input from R.B. D.L. and R.B. wrote the paper.

## References

1. Arendt, D. et al. The origin and evolution of cell types. Nat. Rev. Genet. 17, 744–757 (2016).

2. Zeng, H. What is a cell type and how to define it? Cell 185, 2739–2755 (2022).

3. Wilton, D. K., Dissing-Olesen, L. & Stevens, B. Neuron-Glia Signaling in Synapse Elimination. Annu. Rev. Neurosci. 42, 107–127 (2019).

4. Kofuji, P. & Araque, A. Astrocytes and Behavior. Annu. Rev. Neurosci. 44, 49–67 (2021).

5. Roberts, R. J. V., Pop, S. & Prieto-Godino, L. L. Evolution of central neural circuits: state of the art and perspectives. Nat. Rev. Neurosci. 23, 725–743 (2022).

6. Tosches, M. A. Developmental and genetic mechanisms of neural circuit evolution. Dev. Biol. 431, 16– 25 (2017).

7. Barsotti, E., Correia, A. & Cardona, A. Neural architectures in the light of comparative connectomics. Curr. Opin. Neurobiol. 71, 139–149 (2021).

8. Herculano-Houzel, S. Life history changes accompany increased numbers of cortical neurons: A new framework for understanding human brain evolution. Prog. Brain Res. 250, 179–216 (2019).

9. Kebschull, J. M. et al. Cerebellar nuclei evolved by repeatedly duplicating a conserved cell-type set. Science 370, (2020).

10. Hain, D. et al. Molecular diversity and evolution of neuron types in the amniote brain. Science 377, eabp8202 (2022).

11. Tosches, M. A. et al. Evolution of pallium, hippocampus, and cortical cell types revealed by single-cell transcriptomics in reptiles. Science 360, 881–888 (2018).

12. Woych, J. et al. Cell-type profiling in salamanders identifies innovations in vertebrate forebrain evolution. Science 377, eabp9186 (2022).

13. Shafer, M. E. R., Sawh, A. N. & Schier, A. F. Gene family evolution underlies cell-type diversification in the hypothalamus of teleosts. Nat Ecol Evol 6, 63–76 (2022).

14. Siletti, K. et al. Transcriptomic diversity of cell types across the adult human brain. Science 382, eadd7046 (2023).

15. Yao, Z., et al. A high-resolution transcriptomic and spatial atlas of cell types in the whole mouse brain. bioRxiv (2023) doi:10.1101/2023.03.06.531121.

16. Herculano-Houzel, S., Mota, B. & Lent, R. Cellular scaling rules for rodent brains. Proc. Natl. Acad. Sci. U. S. A. 103, 12138–12143 (2006).

17. Mu, S. et al. 3D reconstruction of cell nuclei in a full Drosophila brain. bioRxiv 2021.11.04.467197 (2021) doi:10.1101/2021.11.04.467197.

18. Vosshall, L. B. Into the mind of a fly. Nature 450, 193–197 (2007).

19. Venken, K. J. T., Simpson, J. H. & Bellen, H. J. Genetic manipulation of genes and cells in the nervous system of the fruit fly. Neuron 72, 202–230 (2011).

20. Scheffer, L. K. & Meinertzhagen, I. A. The Fly Brain Atlas. Annu. Rev. Cell Dev. Biol. 35, 637–653 (2019).

21. Davie, K. et al. A Single-Cell Transcriptome Atlas of the Aging Drosophila Brain. Cell 174, 982– 998.e20 (2018).

22. Croset, V., Treiber, C. D. & Waddell, S. Cellular diversity in the Drosophila midbrain revealed by single-cell transcriptomics. Elife 7, (2018).

23. Li, H. et al. Fly Cell Atlas: A single-nucleus transcriptomic atlas of the adult fruit fly. Science 375, eabk2432 (2022).

24. Markow, T. A. Host use and host shifts in Drosophila. Curr Opin Insect Sci 31, 139–145 (2019).

25. Auer, T. O. & Benton, R. Sexual circuitry in Drosophila. Curr. Opin. Neurobiol. 38, 18–26 (2016).

26. Garrigan, D. et al. Genome sequencing reveals complex speciation in the Drosophila simulans clade. Genome Res. 22, 1499–1511 (2012).

27. Schrider, D. R., Ayroles, J., Matute, D. R. & Kern, A. D. Supervised machine learning reveals introgressed loci in the genomes of Drosophila simulans and D. sechellia. PLoS Genet. 14, e1007341 (2018).

28. Auer, T. O., Shahandeh, M. P. & Benton, R. Drosophila sechellia: A Genetic Model for Behavioral Evolution and Neuroecology. Annu. Rev. Genet. (2021) doi:10.1146/annurev-genet-071719-020719.

29. Farine, J.-P., Legal, Luc, Moreteau, B. & Le Quere, J.-L. Volatile components of ripe fruits of Morinda citrifolia and their effects on Drosophila. Phytochemistry 41, 433–438 (1996).

30. Matute, D. R. & Ayroles, J. F. Hybridization occurs between Drosophila simulans and D. sechellia in the Seychelles archipelago. J. Evol. Biol. 27, 1057–1068 (2014).

31. Lavista-Llanos, S. et al. Dopamine drives Drosophila sechellia adaptation to its toxic host. Elife 3, (2014).

32. Prieto-Godino, L. L. et al. Evolution of Acid-Sensing Olfactory Circuits in Drosophilids. Neuron 93, 661– 676.e6 (2017).

33. Auer, T. O. et al. Olfactory receptor and circuit evolution promote host specialization. Nature 579, 402– 408 (2020).

34. Álvarez-Ocaña, R. et al. Odor-regulated oviposition behavior in an ecological specialist. Nat. Commun. 14, 3041 (2023).

35. Shahandeh, M. P. et al. Evolution of circadian behavioral plasticity through cis-regulatory divergence of a neuropeptide gene. bioRxiv 2023.07.05.547553 (2023) doi:10.1101/2023.07.05.547553.

36. Reisenman, C. E., Wong, J., Vedagarbha, N., Livelo, C. & Scott, K. Taste adaptations associated with host specialization in the specialist Drosophila sechellia. J. Exp. Biol. 226, (2023).

37. Dey, M., Brown, E., Charlu, S., Keene, A. & Dahanukar, A. Evolution of fatty acid taste in drosophilids. Cell Rep. 42, 113297 (2023).

38. Dekker, T., Ibba, I., Siju, K. P., Stensmyr, M. C. & Hansson, B. S. Olfactory shifts parallel superspecialism for toxic fruit in Drosophila melanogaster sibling, D. sechellia. Curr. Biol. 16, 101–109 (2006).

39. Ibba, I., Angioy, A. M., Hansson, B. S. & Dekker, T. Macroglomeruli for fruit odors change blend preference in Drosophila. Sci. Nat. 97, 1059–1066 (2010).

40. Takagi, S. et al. Sensory neuron population expansion enhances odour tracking through relaxed projection neuron adaptation. bioRxiv 2023.09.15.556782 (2023) doi:10.1101/2023.09.15.556782.

41. Zheng, G. X. Y. et al. Massively parallel digital transcriptional profiling of single cells. Nat. Commun. 8, 14049 (2017).

42. Deng, B. et al. Chemoconnectomics: Mapping Chemical Transmission in Drosophila. Neuron 101, 876–893.e4 (2019).

43. Bates, A. S. et al. The natverse, a versatile toolbox for combining and analysing neuroanatomical data. Elife 9, (2020).

44. Hobert, O. A map of terminal regulators of neuronal identity in Caenorhabditis elegans. Wiley Interdiscip. Rev. Dev. Biol. 5, 474–498 (2016).

45. Özel, M. N. et al. Coordinated control of neuronal differentiation and wiring by sustained transcription factors. Science 378, eadd1884 (2022).

46. Bai, L., Goldman, A. L. & Carlson, J. R. Positive and negative regulation of odor receptor gene choice in Drosophila by acj6. J. Neurosci. 29, 12940–12947 (2009).

47. Jafari, S. et al. Combinatorial activation and repression by seven transcription factors specify Drosophila odorant receptor expression. PLoS Biol. 10, e1001280 (2012).

48. Kurusu, M. et al. Genetic control of development of the mushroom bodies, the associative learning centers in the Drosophila brain, by the eyeless, twin of eyeless, and Dachshund genes. Proc. Natl. Acad. Sci. U. S. A. 97, 2140–2144 (2000).

49. Guo, X., Zhang, Y., Huang, H. & Xi, R. A hierarchical transcription factor cascade regulates enteroendocrine cell diversity and plasticity in Drosophila. Nat. Commun. 13, 6525 (2022).

50. Freeman, M. R. Drosophila Central Nervous System Glia. Cold Spring Harb. Perspect. Biol. 7, (2015).

51. Cavigliasso, F. et al. Cis-regulatory polymorphism at fiz ecdysone oxidase contributes to polygenic adaptation to malnutrition in Drosophila. bioRxiv 2023.08.28.555138 (2023) doi:10.1101/2023.08.28.555138.

52. Dworkin, I. & Jones, C. D. Genetic changes accompanying the evolution of host specialization in Drosophila sechellia. Genetics 181, 721–736 (2009).

53. Volkenhoff, A. et al. Glial Glycolysis Is Essential for Neuronal Survival in Drosophila. Cell Metab. 22, 437–447 (2015).

54. Watanabe, K. et al. Interspecies Comparative Analyses Reveal Distinct Carbohydrate-Responsive Systems among Drosophila Species. Cell Rep. 28, 2594–2607.e7 (2019).

55. Day, E. H., Hua, X. & Bromham, L. Is specialization an evolutionary dead end? Testing for differences in speciation, extinction and trait transition rates across diverse phylogenies of specialists and generalists. J. Evol. Biol. 29, 1257–1267 (2016).

56. Nässel, D. R., Enell, L. E., Santos, J. G., Wegener, C. & Johard, H. A. D. A large population of diverse neurons in the Drosophila central nervous system expresses short neuropeptide F, suggesting multiple distributed peptide functions. BMC Neurosci. 9, 90 (2008).

57. Nässel, D. R. & Wegener, C. A comparative review of short and long neuropeptide F signaling in invertebrates: Any similarities to vertebrate neuropeptide Y signaling? Peptides 32, 1335–1355 (2011).

58. Fadda, M. et al. Regulation of Feeding and Metabolism by Neuropeptide F and Short Neuropeptide F in Invertebrates. Front. Endocrinol. 10, 64 (2019).

59. Bittern, J. et al. Neuron-glia interaction in the Drosophila nervous system. Dev. Neurobiol. 81, 438–452 (2021).

60. Rifkin, S. A., Kim, J. & White, K. P. Evolution of gene expression in the Drosophila melanogaster subgroup. Nat. Genet. 33, 138–144 (2003).

61. Khaitovich, P. et al. A neutral model of transcriptome evolution. PLoS Biol. 2, E132 (2004).

62. Nogueira Alves, A., Oliveira, M. M., Koyama, T., Shingleton, A. & Mirth, C. K. Ecdysone coordinates plastic growth with robust pattern in the developing wing. Elife 11, (2022).

63. Turner, B. M. Epigenetic responses to environmental change and their evolutionary implications. Philos. Trans. R. Soc. Lond. B Biol. Sci. 364, 3403–3418 (2009).

64. Young, M. D. & Behjati, S. SoupX removes ambient RNA contamination from droplet-based single-cell RNA sequencing data. Gigascience 9, giaa151 (2020).

65. Hafemeister, C. & Satija, R. Normalization and variance stabilization of single-cell RNA-seq data using regularized negative binomial regression. Genome Biol. 20, 296 (2019).

66. McGinnis, C. S., Murrow, L. M. & Gartner, Z. J. DoubletFinder: Doublet Detection in Single-Cell RNA Sequencing Data Using Artificial Nearest Neighbors. Cell Syst 8, 329–337.e4 (2019).

67. Emms, D. M. & Kelly, S. OrthoFinder: phylogenetic orthology inference for comparative genomics. Genome Biol. 20, 238 (2019).

68. Hao, Y. et al. Integrated analysis of multimodal single-cell data. Cell 184, 3573–3587.e29 (2021).

69. Chen, E. Y. et al. Enrichr: interactive and collaborative HTML5 gene list enrichment analysis tool. BMC Bioinformatics 14, 128 (2013).

70. Kuleshov, M. V. et al. Enrichr: a comprehensive gene set enrichment analysis web server 2016 update. Nucleic Acids Res. 44, W90–7 (2016).

71. Ostrovsky AD, Goetz L, Jefferis GSXE. Drosophila melanogaster template brains. (2014) doi:10.5281/zenodo.10591.

72. Ostrovsky AD, Goetz L, Jefferis GSXE. Drosophila simulans template brains. (2014) doi:10.5281/zenodo.10594.

73. Jefferis GSXE, Benton R. DsecF Drosophila sechellia Female Template Brain. (2019) doi:10.5281/zenodo.2562141.

74. Winkler, B. et al. Brain inflammation triggers macrophage invasion across the blood-brain barrier in Drosophila during pupal stages. Sci Adv 7, eabh0050 (2021).

