## Supplementary Information for "Comparative single-cell transcriptomic atlases reveal conserved and divergent features of drosophilid central brains"

**Supplementary Table 1. Summary of the metrics from the single-nucleus RNA-sequencing (snRNA-seq) experiments**

**Supplementary Table 2. Marker genes for the 48 annotated and 16 unannotated cell types**

**Supplementary Table 3. 368 broadly expressed genes (>5% of central brain cells in *D. melanogaster*) with conserved expression patterns (Spearman’s ρ > 0.7)**

**Supplementary Table 4. 896 genes with specific expression (<10 of the 48 annotated cell types) and conserved expression patterns (expressed in >30% of cells across all three species)**

The subset encoding predicted/known transcription factors (TFs) are shown separately on the second worksheet.

**Supplementary Table 5. Differentially expressed genes identified from pairwise comparisons of three drosophilid species in a cell type-specific manner**

**Supplementary Table 6. 80 genes with putative expression changes specific to the *D. sechellia* lineage**

**Supplementary Figure 1. Integrated single-cell transcriptomic atlases of *D. melanogaster*, *D. simulans* and *D. sechellia***

(a,b) tSNE (a) and UMAP (b) plots of the integrated dataset after RPCA integration of single-cell transcriptomes of *D. melanogaster*, *D. simulans* and *D. sechellia* central brain cells. Cells are colored by the SNN-based clustering, and clusters are numbered according to their sizes, starting with the largest as 0.

(c,d) tSNE (c) and UMAP (d) plots with cells colored by the expression of *pros* or *Imp*.

(e,f) tSNE (e) and UMAP (f) plots with cells colored by their predicted molecular identities. Neuronal cells are identified through markers for neurotransmitter production, color-coded according to GABAergic (*Gad1*), monoaminergic (*Vmat*), glutamatergic (*Vglut*) and cholinergic (*VAChT*) designations. Glial cells, labeled by *repo* expression, are depicted in black. Cells failing to meet expression thresholds for these markers appear in gray.

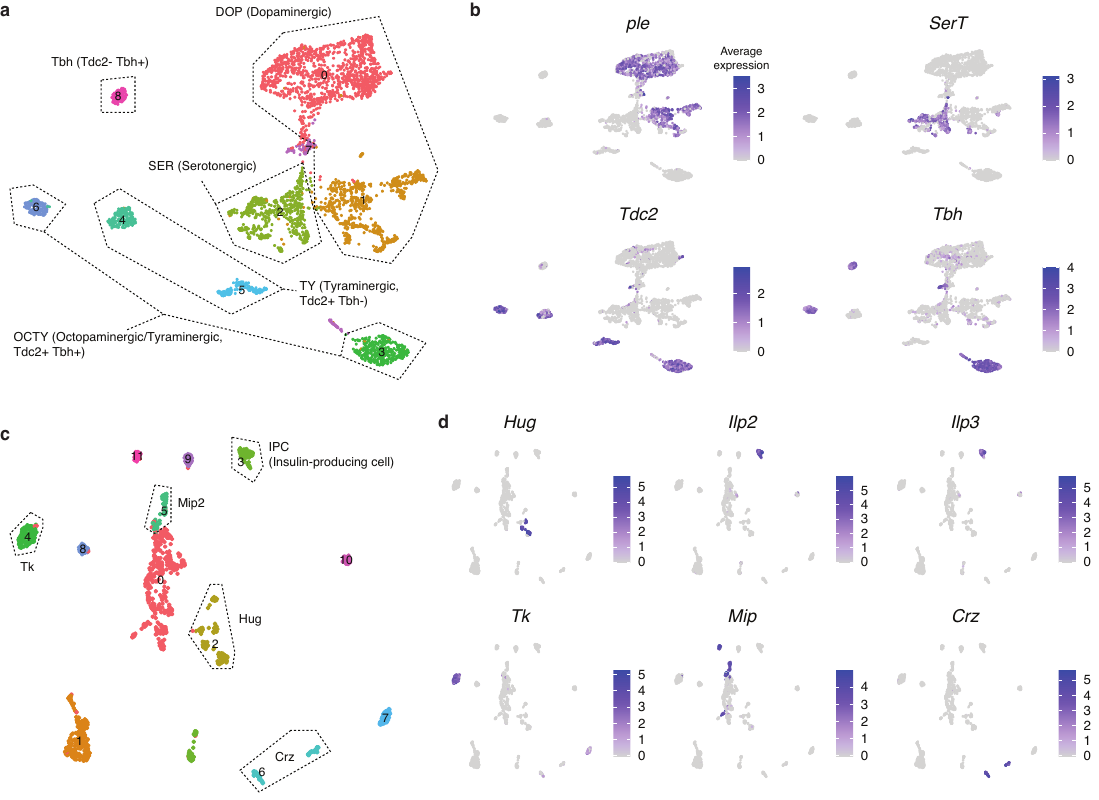

**Supplementary Figure 2. Annotation of cell types in subclusters within monoaminergic and peptidergic cell clusters**

(a) UMAP plot displaying subclusters of a monoaminergic cluster. Cells are colored by their SNN-based subclustering, and labeled according to their molecular identity inferred from marker gene expression as shown in (b).

(b) UMAP plots showing the expression of marker genes for subclusters of a monoaminergic cluster, including dopaminergic (*ple*), serotonergic (*SerT*), tyraminergic (*Tdc2*) and octopaminergic (*Tbh*) subclusters.

(c) UMAP plot displaying subclusters of a peptidergic cluster. Cells are colored by their SNN-based subclustering, and labeled according to their molecular identity inferred from marker gene expression as shown in (d).

(d) UMAP plots showing the expression of marker genes for subclusters of a peptidergic cluster, including subclusters producing hugin (*Hug+*), insulin (*Ilp2+* and *Ilp3+*), tachykinin (*Tk+*), MIP (*Mip+*) and corazonin (*Crz+*) subclusters.

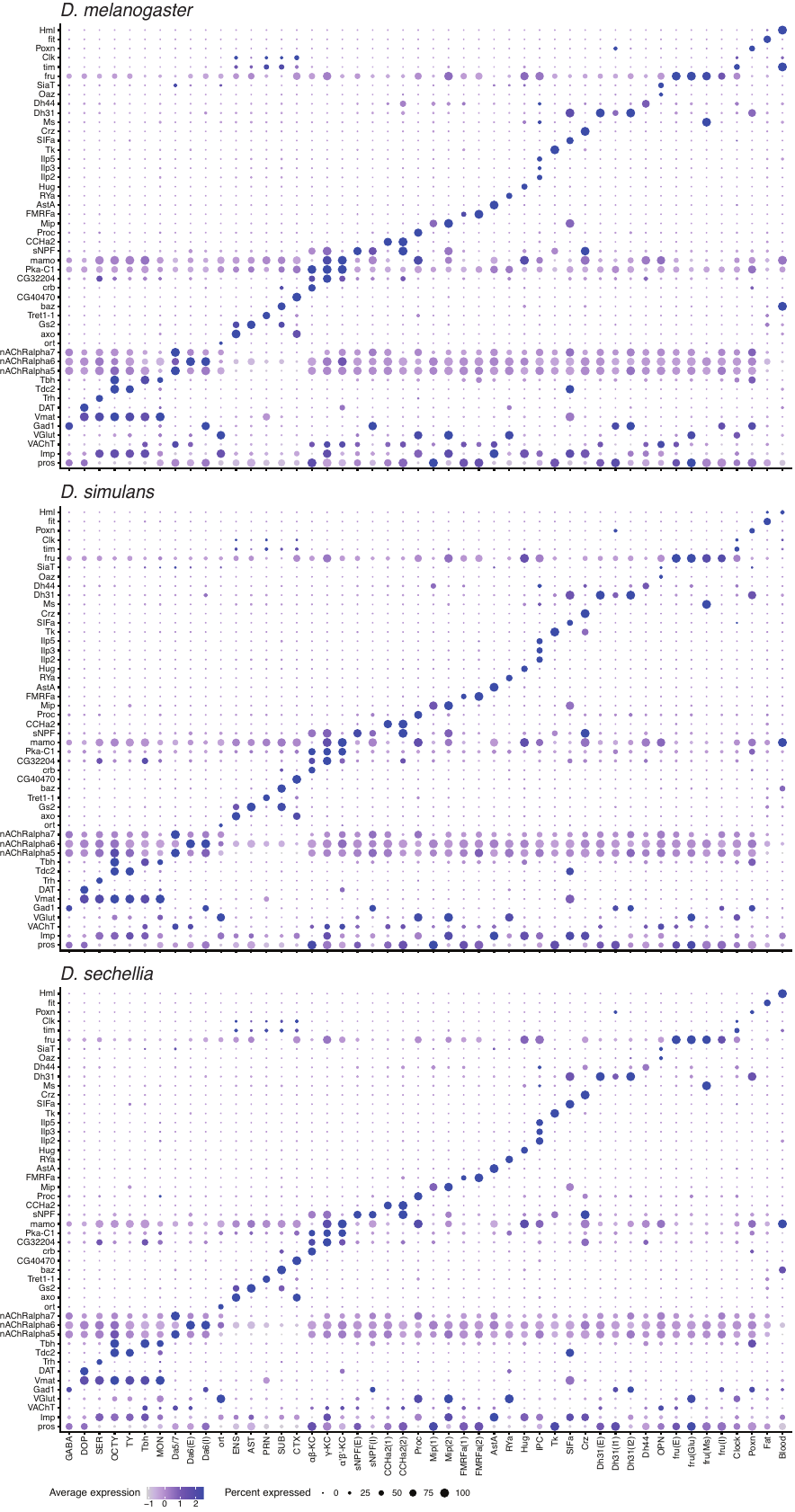

**Supplementary Figure 3. Conserved cell type marker expression across *D. melanogaster*, *D. simulans* and *D. sechellia***

Dot plots summarizing the expression of marker genes used for the annotation of 48 cell types in the drosophilid species.

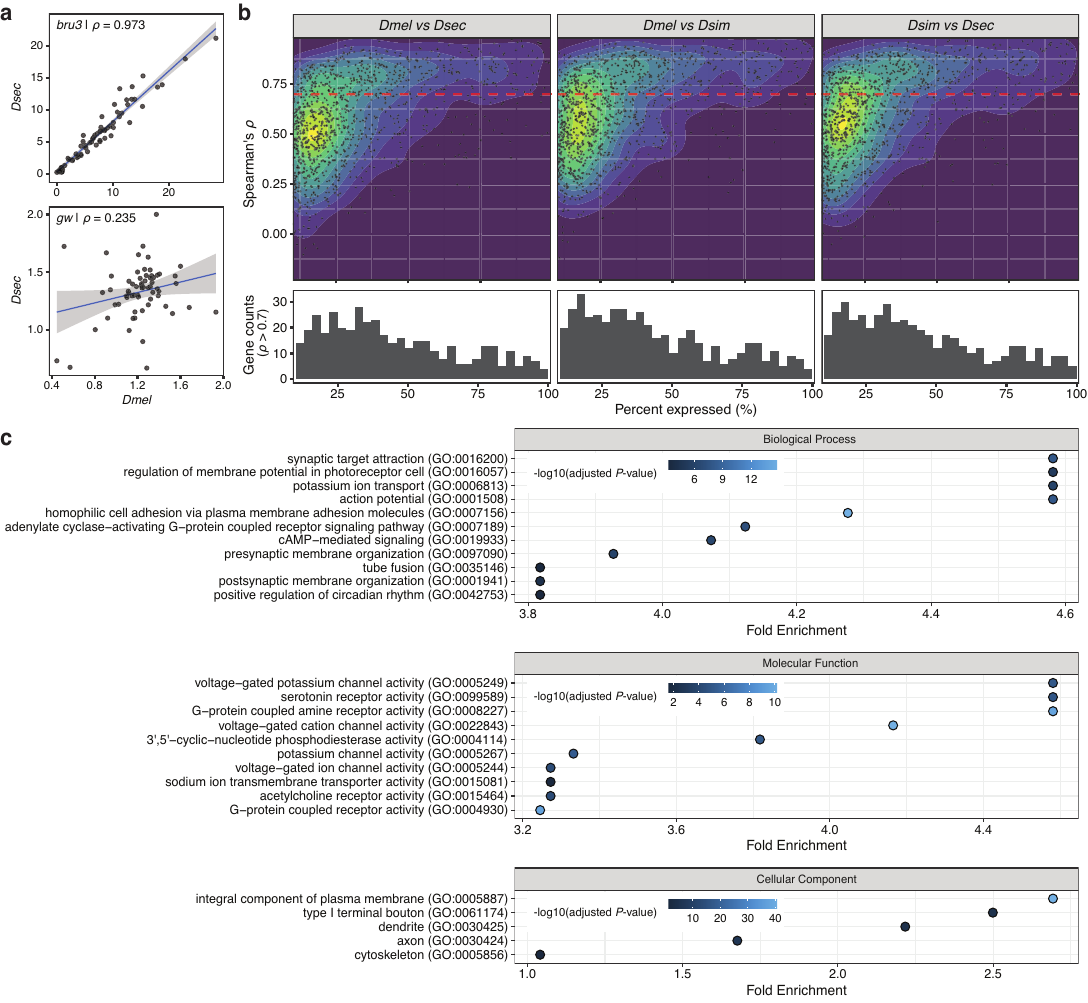

**Supplementary Figure 4. Genes with conserved expression patterns across the central brains of *D. melanogaster*, *D. simulans* and *D. sechellia***

(a) Scatter plots illustrating the expression pattern comparisons of two genes, *bru3* (top) and *gw* (bottom), between *D. melanogaster* and *D. sechellia*. Each point corresponds to one of the 64 cell types. Average expression levels of *D. melanogaster* are shown on the x-axis, and *D. sechellia* expression levels are shown on the y-axis. Smoothed lines depict generalized linear model fits.

(b) Top: two-dimensional density plots of gene expression pattern similarity for three pairwise comparisons. Each point in the plot corresponds to one of 1,686 analyzed genes. The y-axis shows the expression pattern similarity, which is estimated by the correlation value (Spearman's *ρ*). Bottom: histograms of gene count with highly conserved expression patterns (*ρ* > 0.7) are shown. The x-axis shows the percentage of central brain cells expressing each gene.

(c) GO analysis for 368 genes with highly conserved expression patterns (*ρ* > 0.7) across all three pairwise comparisons.

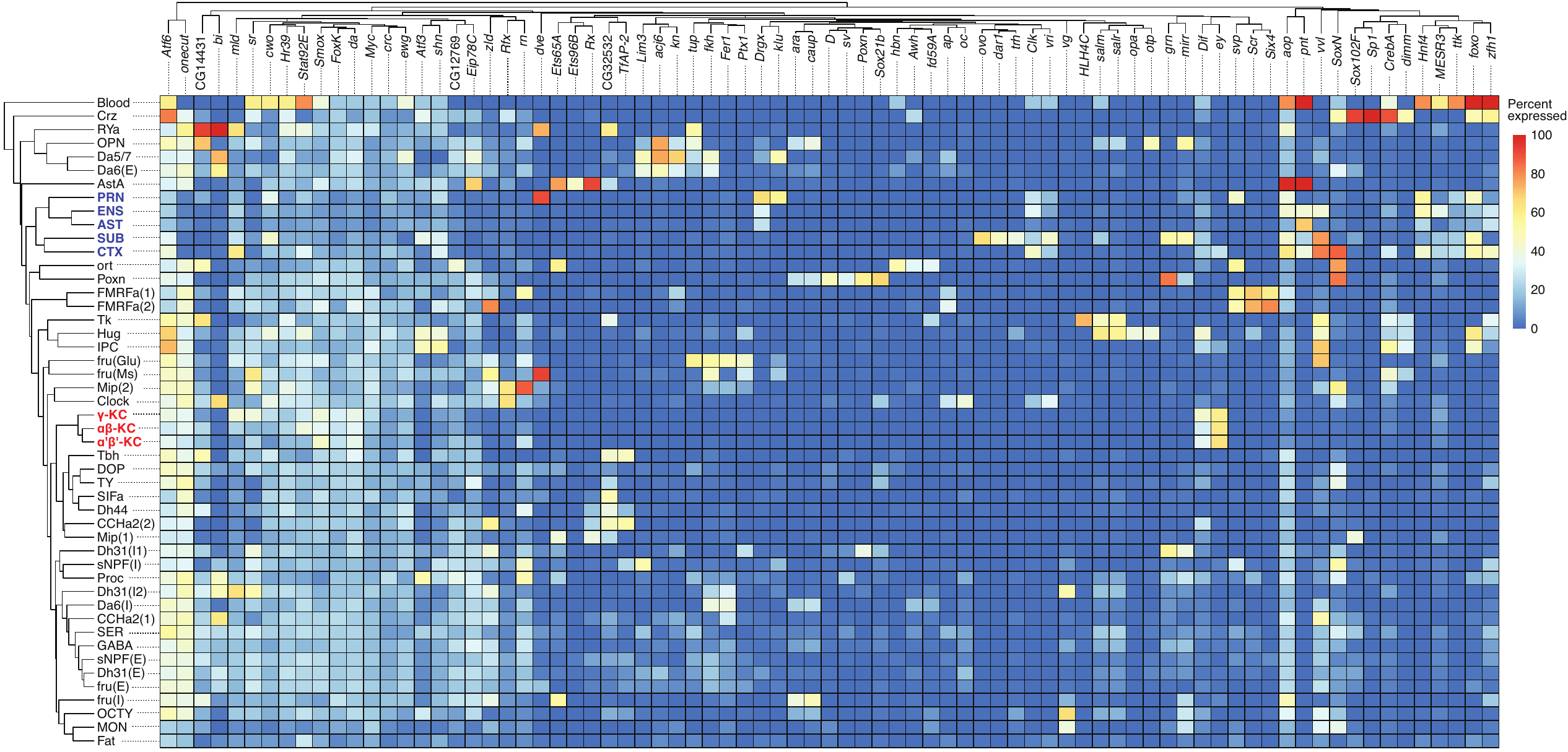

**Supplementary Figure 5. Transcription factor fingerprints for annotated cell types**

A heatmap illustrating the percentage of cells expressing 79 transcription factor genes across 48 annotated cell types, derived from the *D. melanogaster* dataset. Both genes and cell types are clustered through hierarchical clustering. Labels for glial cell types are colored blue and Kenyon cell types are colored red.

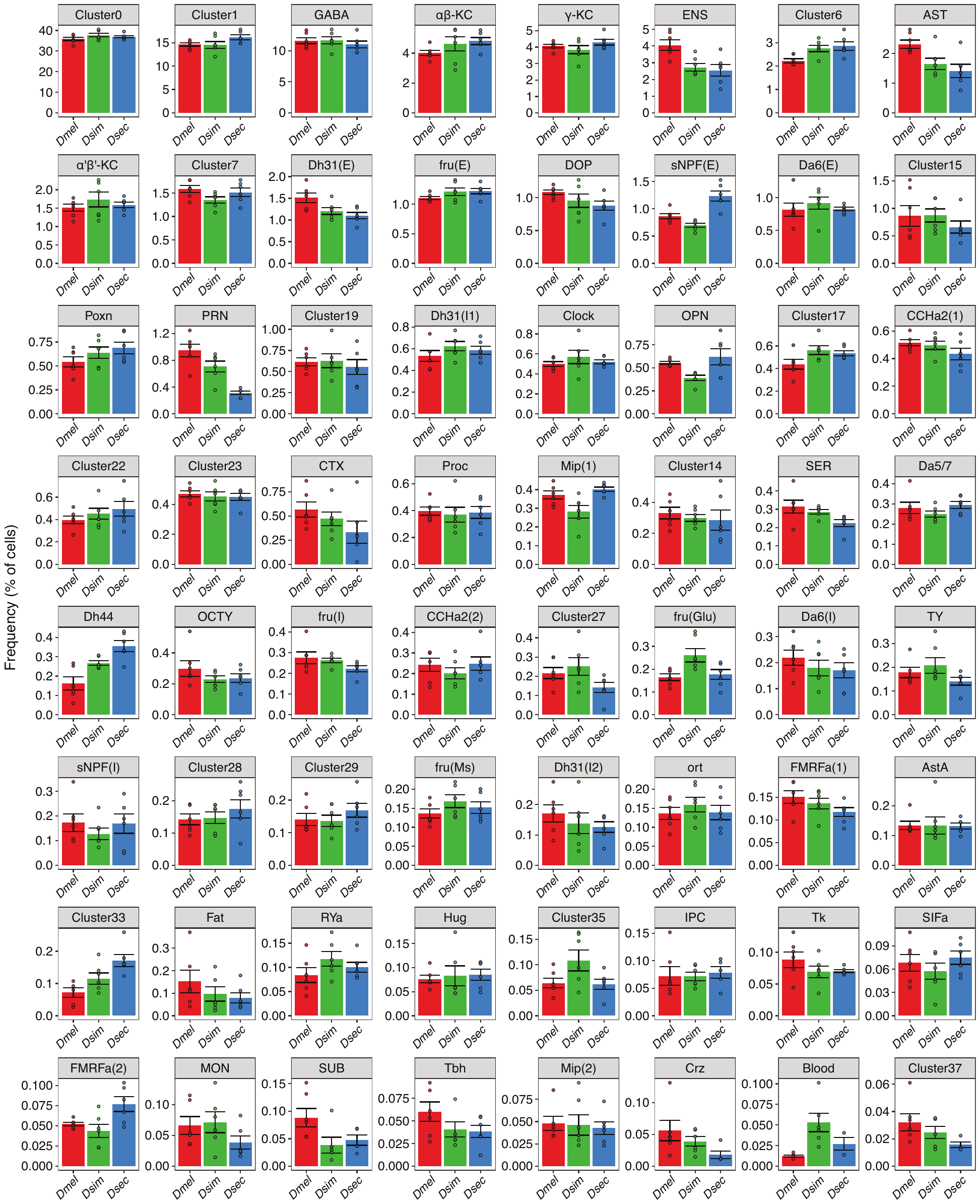

**Supplementary Figure 6. Frequency comparisons across homologous cell types**

Bar plots showing comparisons of cell type frequencies of 64 annotated and unannotated cell types across the drosophilid species. Each point corresponds to one of six replicates.

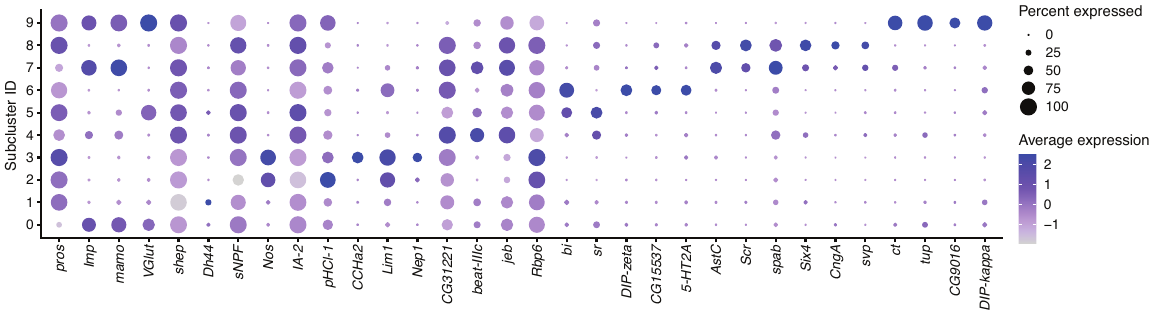

**Supplementary Figure 7. Interspecific subtype frequency variation in sNPF(E) cell types**

A dot plot summarizing the expression of marker genes for the top ten largest subclusters of sNPF(E) cell types.

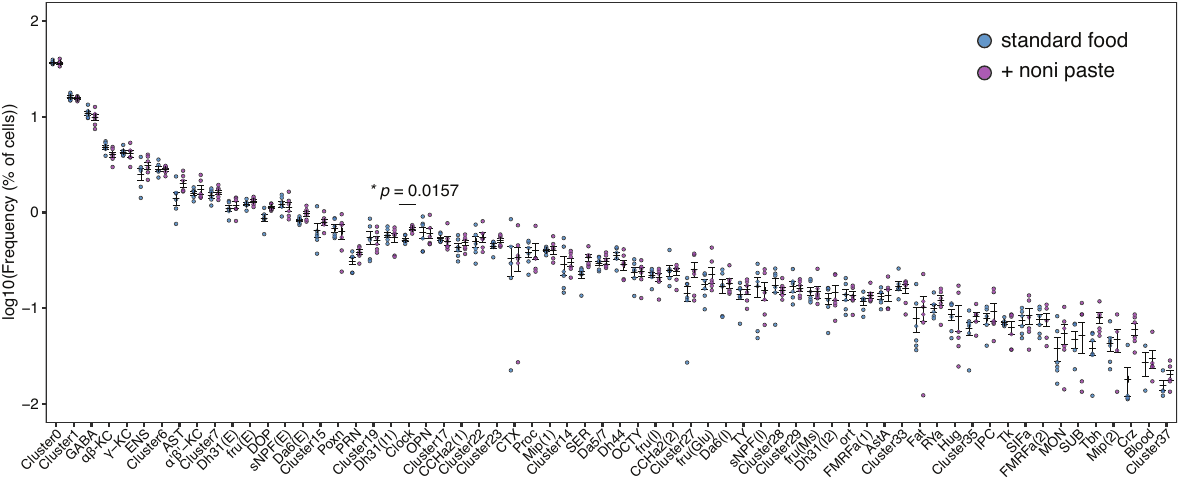

**Supplementary Figure 8. Impact of noni paste supplement on cell type frequencies in the *D. sechellia* central brain**

A dot plot showing comparisons of brain cell type frequencies between *D. sechellia* grown in standard medium and noni paste-supplemented medium. Cell types are shown on the x-axis and frequencies are shown on the y-axis. Each point corresponds to one of six replicates. The error bars represent the standard error of the mean (SEM). A paired t-test was conducted to assess statistical significance, with *p*-values adjusted using false discovery rate (FDR).

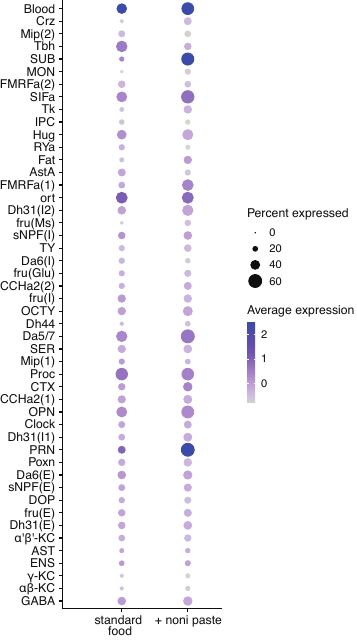

**Supplementary Figure 9. Expression changes of CG5151 in *D. sechellia* upon noni-paste supplement**

Dot plots summarizing the expression changes of CG5151 across 48 annotated cell types of brains of *D. sechellia* grown on standard food or with noni paste supplement. In the latter condition, CG5151 is upregulated in PRN and SUB, but this increase is statistically significant only in PRN.
